# *longfin* causes *cis*-ectopic expression of the *kcnh2a ether-a-go-go* K^+^ channel to autonomously prolong fin outgrowth

**DOI:** 10.1101/790329

**Authors:** Scott Stewart, Heather K. Le Bleu, Gabriel A. Yette, Astra L. Henner, Amy E. Robbins, Joshua A. Braunstein, Kryn Stankunas

## Abstract

Organs stop growing to achieve a characteristic size and shape in scale with the animal’s body. Likewise, regenerating organs sense injury extents to instruct appropriate replacement growth. Fish fins exemplify both phenomena through their tremendous diversity of form and remarkably robust regeneration. The classic zebrafish mutant *longfin^t2^* develops and regenerates dramatically elongated fins and underlying ray skeleton. We show *longfin^t2^* chromosome 2 overexpresses the *ether-a-go-go*-related voltage-gated potassium channel *kcnh2a.* Genetic disruption of *kcnh2a in cis* rescues *longfin^t2^,* indicating *longfin^t2^* is a regulatory *kcnh2a* allele. We find *longfin^t2^* fin overgrowth originates from prolonged outgrowth periods including by showing Kcnh2a chemical inhibition during late stage regeneration fully suppresses overgrowth. Cell transplantations demonstrate *longfin^t2^*-ectopic *kcnh2a* acts tissue autonomously within the fin intra-ray mesenchymal lineage. Temporal inhibition of the Ca^2+^-dependent phosphatase calcineurin indicates it likewise entirely acts late in regeneration to attenuate fin outgrowth. Epistasis experiments suggest *longfin^t2^*-expressed Kcnh2a inhibits calcineurin output to supersede growth cessation signals. We conclude ion signaling within the growth-determining mesenchyme lineage controls fin size by tuning outgrowth periods rather than altering positional information or cell-level growth potency.

## INTRODUCTION

Animal organs develop to and maintain a species-specific size and shape in scale with the individual’s overall body. Remarkably, regenerating organs can sense injury extents to restore their original proportions. Fish fins neatly demonstrate both organ size establishment and regeneration mysteries. Fins display tremendous morphological diversity to optimize swimming, predator avoidance, and courtship, while also contributing to fishes’ aesthetic appeal (Nelson et al., 2016). Further, most fins, including those of the zebrafish model organism, regenerate to their original form, providing a striking example of organ size control and scaling.

Zebrafish fin skeletons comprised of segmented, cylindrical bony rays define the overall fin form. Each ray segment embodies two opposing hemi-cylindrical bones produced by tightly associated osteoblasts. A stratified epidermis lines and vasculature, sensory axons, and fibroblasts reside within rays. During larval development, fins initiate rapid asynchronous allometric growth culminating in their mature forms, including the familiar bi-lobed shape of the well-studied caudal fin (Goldsmith et al., 2006). Adult fins switch to isometric growth, slowly expanding in scale with the rest of the body throughout the animal’s life. Adult fin regeneration engages the appropriate extent of rapid allometric growth to restore a properly proportioned appendage (Goldsmith et al., 2006; Iovine and Johnson, 2000).

The zebrafish fin regeneration model affords a tractable platform to determine fin size control mechanisms. No matter where a fin is amputated, the regenerated fin restores its original size and shape (Chen and Poss, 2017; Sehring and Weidinger, 2020). Fin amputation triggers a wound epithelium formed by migrating epidermal cells. De-differentiation of mature cells to lineage-restricted progenitors then generates a regenerative blastema for each ray (Knopf et al., 2011; Singh et al., 2012; Sousa et al., 2011; Stewart and Stankunas, 2012; Tu and Johnson, 2011). Heterogeneous blastemas organize by cell lineage and state to enable progressive regeneration (reviewed in (Wehner and Weidinger, 2015)). De-differentiated, proliferative osteoblasts (pObs) migrate to blastema peripheries and hierarchically arrange along the distal-to-proximal axis of the blastema, with the most progenitor “state” cells distally concentrated (Stewart et al., 2014). Similarly, fibroblasts residing between hemi-rays de-differentiate, form the major blastema mesenchyme population radially interior to pObs, and then contribute to regenerated intra-ray tissue (Tornini et al., 2016). An outgrowth phase follows that integrates spatially segregated proliferation (distal) and differentiation (proximal) activities for the progressive restoration of replacement tissue.

Developmental signaling pathways promote tissue interactions and cell behaviors to replace lost cells and re-form mature, patterned tissue (Wehner and Weidinger, 2015). A distal blastema “organizing center” produces essential Wnt signals that coordinates other pathways including FGFs and BMPs (Wehner et al., 2014). FGFs likely are the primary mitogens (Lee et al., 2005; Lee et al., 2009; Poss et al., 2000; Shibata et al., 2016), whereas BMP signaling is implicated in osteoblast differentiation (Quint et al., 2002; Smith et al., 2006; Stewart et al., 2014). Yet, cell behaviors and their mechanistic control that ensure cessation of regrowth once the proper fin size is reached are unresolved. Prevailing models for fin size restoration suggest positional identities of cells at an injury site establish appropriate outgrowth-determining blastema pre-patterns (Lee et al., 2005; Rabinowitz et al., 2017; Tornini et al., 2016). The nature of such cellular memories, how they establish blastema positional information, and how those pre-patterns are converted into outgrowth extents all remain largely mysterious.

Genetic studies of adult viable zebrafish mutants with abnormally sized fins provide an entry for molecular insights into fin growth control (Van Eeden et al., 1996). These studies universally implicate ion signaling as a major determinant of fin size. Gain-of-function mutations in the K^+^ channel *kcnk5b* in *another longfin* (*alf*) develop and regenerate long fins (Perathoner et al., 2014). Further, fin overgrowth in the *schleier* mutant is caused by loss of *kcc4a/ slc12a7a*, a K^+^ Cl^-^ cotransporter (Lanni et al., 2019). Ectopic expression of *kcnj13*, coding for a K^+^ channel, caused by a viral insertion or transgene-mediated expression of the Kcnj1b, Kcnj10a, and Kcnk9c K^+^ channels also causes overgrowth (Silic et al., 2020). Finally, loss of *gap junction protein alpha 1b* (*gja1*, also known as *connexin 43*) in *shortfin* mutants suggests involvement of intercellular ion exchange (Iovine and Johnson, 2000). This literature, together with studies in other animals, points to central roles of ion signaling, or “bioelectricity” in organ size establishment (McLaughlin and Levin, 2018). However, how ion signaling dynamics directly alter cell states and behaviors to instruct and/or enable scaled growth is poorly understood.

Cooperating factors that promote and effect ion signaling dynamics for fin size control are unresolved. As one key insight, small molecule inhibition of the Ca^2+^-dependent phosphatase calcineurin causes dramatic fin overgrowth (Kujawski et al., 2014). As a decoder of the ubiquitous cytosolic Ca^2+^ second messenger, calcineurin links ion signaling and protein effector phosphorylation (Crabtree, 1999). Calcineurin may act directly upstream of Kcnk5b (*alf*) to temper outgrowth and restore fin proportions (Daane et al., 2018; Yi et al., 2020). In mammals, diverse stimuli activate calcineurin by elevating intracellular Ca^2+^, including antigen engagement of the T-cell receptor complex (Rao, 2009), depolarization of cardiomyocytes (Parra and Rothermel, 2017), and neuronal ion channel activity (Baumgärtel and Mansuy, 2012). Calcineurin dephosphorylates multiple proteins, including ion channels and transcription factors, to promote context-dependent cell behaviors. Characterizing calcineurin’s upstream regulators and downstream effectors in the fin is a promising path to reveal organ size control mechanisms.

The classic dominant *longfin^t2^* (*lof^t2^)* mutant also develops and regenerates exceptionally long fins, producing a “schleierform”, or flowing veil-like appearance (Elias, 1984; Van Eeden et al., 1996) (Fig. 1A and B). Notably, the widely used stock strain *Tüpfel-Longfin* (*TL*) contains compound *lof^t2/t2^; leopard^t1/t1^* (*leo*) mutations, whose respective fin and pigmentation phenotypes allow visual identification of mixed genotypes (Haffter et al., 1996). Developing fins of *lof^t2^* fish do not have a greater maximal growth rate (Iovine and Johnson, 2000). Rather, *lof^t2^* fins fail to return to isometric growth as animals reach maturity (Goldsmith et al., 2006; Iovine and Johnson, 2000). Unlike *kcnk5b* (*alf*) and *kcc4a/ slc12a7a* (*schlier*) mutants as well as calcineurin inhibition, *lof^t2^* overgrown fins lack other defects such as tissue hyper-vascularization and elongated ray segments (Kujawski et al., 2014; Lanni et al., 2019; McMillan et al., 2018; Perathoner et al., 2014; Silic et al., 2020). The latter phenotype likely reflects the independent disruption of joint formation rather than overgrowth since ray segment number and fin size are not correlated; *evx1* mutants, devoid of all fin joints, have normal fin sizes (Schulte et al., 2011; Ton and Iovine, 2013). Regardless, *lof^t2^* provides a uniquely clean model to explore fin size control.

**Figure 1.**
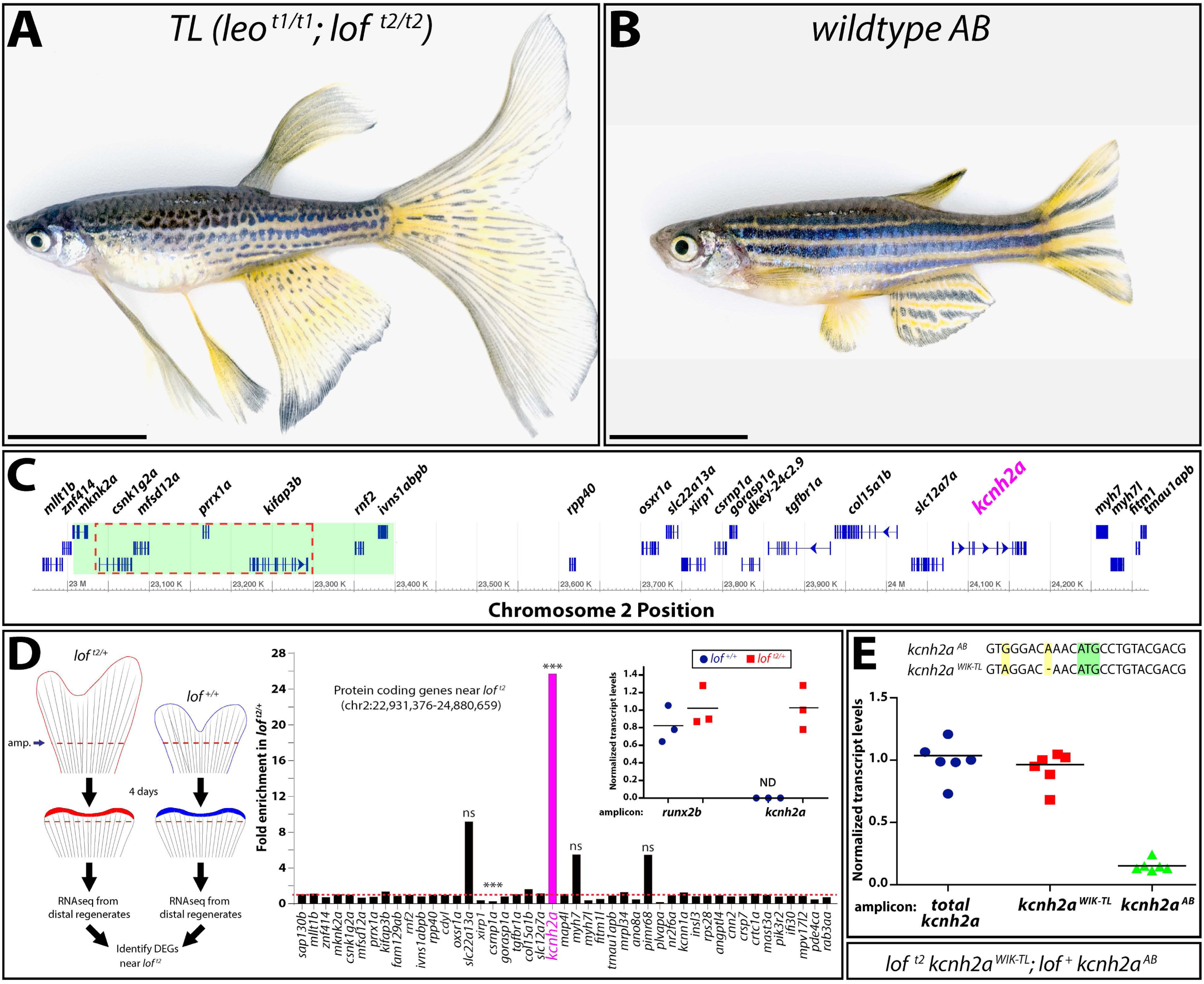
The *lof^t2^* mutation causes *cis*-ectopic expression of *kcnh2a*. **(A**, **B)** Brightfield images of adult *TL (Tüpfel longfin: leo^t1/t1^; lof^t2/t2^)* and wildtype *AB* zebrafish. The scale bar represents 1 cm. (**C)** Schematic diagram of 1.5 Mb region of zebrafish chromosome 2 (chr2). The putative location of the *lof^t2^* mutation is outlined with a dashed red line and the region deleted in the suppressed *lof^jg1^* is highlighted by a green box as determined in (Iovine and Johnson, 2002). **(D) Left.** Schematic of RNA-Seq experimental design to identify genes mis-expressed in regenerated *lof^t2^*. **Right.** RNA-Seq data showing gene expression at chr2:22,931,376-24,880,659 from *lof^t2/+^* relative to clutchmate *lof^+/+^* controls. The asterisks indicate the two differentially expressed genes (p < 10^-4^). ns: not significant. **Inset.** Expression of *kcnh2a* in *lof^t2^***^/+^** (in red) and *lof^+/+^* (in blue) clutchmates determined by RT-qPCR using 4 dpa fin cDNA. Data were normalized to *rpl8* reference expression levels and presented as fold change relative to *lof^t2/+^*. *runx2b* expression levels are shown for comparative purposes and were not significantly changed between the two genotypes. Expression of *kcnh2a* was below limits of detection in *lof^+/+^* fish (indeterminate, ND). Each point represents a cohort of 3 animals. **(E)** RT-qPCR on 4 dpa caudal fin cDNA from *lof^t2^ kcnh2a^WIK-TL^; lof^+^ kcnh2a^AB^* fish to detect chromosome-specific expression *kcnh2a.* Sequences of non-coding *kcnh2a* polymorphisms that specifically amplify either *kcnh2a^WIK-TL^*, located on the *lof^t2^* mutant chr2 (red squares), or *kcnh2a^AB^*, located on *AB* chr2 (green triangles). Data were normalized to total *kcnh2a* (blue circle) levels determined using primers that amplify both alleles indiscriminately. Each data point represents data from an individual fish. RT-qPCR statistical analyses used one-way ANOVA with Tukey’s multiple comparisons tests.

Further motivated by the historical significance and prominence of *lof^t2^*, we sought to identify the cause of its eponymous phenotype. We show the dominant *lof^t2^* fin overgrowth phenotype is caused by *cis* ectopic expression of *kcnh2a*, a K^+^ channel-encoding gene mapping near the *lof* locus. During regeneration, the activity of ectopic Kcnh2a, an ortholog of cardiac arrhythmia-associated *ether-a-go-go* channels, promotes fin overgrowth exclusively by extending the outgrowth period. Labeled *lof^t2^* genetic chimeras demonstrate Kcnh2a acts tissue autonomously within size-determining intra-ray mesenchymal lineage cells. Finally, we provide evidence Kcnh2a disrupts a Ca^2+/^calcineurin pathway that gradually terminates allometric fin outgrowth. Our results suggest readily tunable ion signaling alters organ size by modulating growth periods rather than establishing growth-defining positional information or directly impacting cell cycling rates.

## RESULTS

### *cis*-overexpression of *kcnh2a* in *lof^t2^*

The dominant *lof^t2^* allele that specifically causes dramatic fin overgrowth maps to an essential ∼250 kilobase region (Fig. 1A-C) of chromosome 2 (chr2) (Iovine and Johnson, 2002). Therefore, *lof^t2^* likely results from a dominant negative or gain-of-function mutation rather than haploinsufficiency. One explanation is that *lof^t2^* alters a transcriptional regulatory element causing over-or ectopic expression of a gene(s) within or near the *lof* region (Fig. 1C). We explored this possibility using mRNA sequencing (RNA-seq) to identify differentially expressed genes (DEGs) from 4 days post-amputation (dpa) caudal fin regenerates (Fig. 1D). We reasoned 4 dpa regenerates would highlight primary transcriptional changes because *lof^t2^* and wildtype fin sizes were indistinguishable at this stage of regeneration (Fig. S1, discussed below). We identified 39 increased and 111 decreased DEGs (+/-2-fold change) in *lof^t2^* fins (Table S1). Confining the DEG analysis to the *lof^t2^* region and surrounding genes (Fig. 1C), a single transcript roughly 1 megabase from *lof^t2^*, *potassium voltage-gated channel, subfamily H (eag-related), member 2a* (*kcnh2a*), was greatly elevated in *lof^t2/+^* animals (+ 25.8 fold) (Fig. 1D). One gene, *csrnp1a*, showed decreased transcript levels (−3.9 fold). Quantitative RT-PCR (RT-qPCR) confirmed *kcnh2a* was expressed uniquely in *lof^t2/+^* fin regenerates, at levels similar to the *runx2b* osteoblast-lineage transcription factor (Fig. 1D inset).

To determine if *lof^t2^* affects *kcnh2a* expression in *cis* or *trans*, we leveraged sequence polymorphisms (Butler et al., 2015) in the 5’ UTR that distinguish *kcnh2a* from *TL* and *WIK* (*kcnh2a^WIK-TL^)* versus our wildtype *AB* (*kcnh2a^AB^*) strains (Fig. 1E). We used allele-specific primers and RT-qPCR to observe that *kcnh2a^WIK-TL^* transcripts accounted for nearly all *kcnh2a* expression in *lof^t2^ kcnh2a^WIK-TL^; lof^+^ kcnh2a^AB^* regenerating caudal fins (Fig. 1E; p < 0.0001). The apparent 10% residual contribution of *kcnh2a^AB^* to total *kcnh2a* expression likely reflects partial primer cross-hybridization. We conclude most, if not all, *kcnh2a* expression in *lof^t2^* regenerating fins arises from *cis* effects of the *lof^t2^* mutation.

### *lof^t2^* is a regulatory neomorph allele of *kcnh2a*

We used CRISPR/Cas9 to mutate *kcnh2a* in the *TL* (*leo^t1/t1^; lof^t2/t2^*) background to determine if overexpressed *kcnh2a* causes *lof^t2^* fin overgrowth. We outcrossed founders to wildtype fish and identified numerous obligatory *lof^t2/+^* F_1_ animals with normal sized fins carrying germline-transmitted *kcnh2a* loss-of-function alleles. One of these, *kcnh2a^b1391^,* contained an 8 bp deletion causing a frameshift at codon 3 of the predicted polypeptide. Homozygous *lof^t2/t2^; kcnh2a^b1391/b1391^* fish were phenotypically normal with fin sizes indistinguishable from wildtype clutchmates (Fig. 2A and B; p < 0.0001 between *lof^t2/t2^* and either wildtype or *lof^t2/t2^*; *kcnh2a^b1391/b1391^* fish; no difference between wildtype and *lof^t2/t2^*; *kcnh2a^b1391/b1391^* genotypes). Since *lof^t2^ kcnh2a^b1391^; lof^+^ kcnh2a^+^* and *lof^t2^ kcnh2a^b1391^; lof^t2^ kcnh2a^+^* fish developed normal and long fins, respectively, the *kcnh2a^b1391^* allele uniquely suppresses *lof^t2^* in *cis*. Regenerative outgrowth also was identical between wildtype and *lof^t2/t2^; kcnh2a^b1391/b1391^* caudal fins (Fig. 2C-F). Finally, *lof^t2^*-linked *kcnh2a* had no exon or intron/exon boundary non-synonymous mutations or unappreciated SNPs. We conclude *cis*-ectopic expression of *kcnh2a* on the *lof^t2^* chr2 causes the dominant *lof^t2^* fin overgrowth phenotype. Therefore, *lof^t2^* likely is a regulatory *kcnh2a* neomorphic allele.

**Figure 2.**
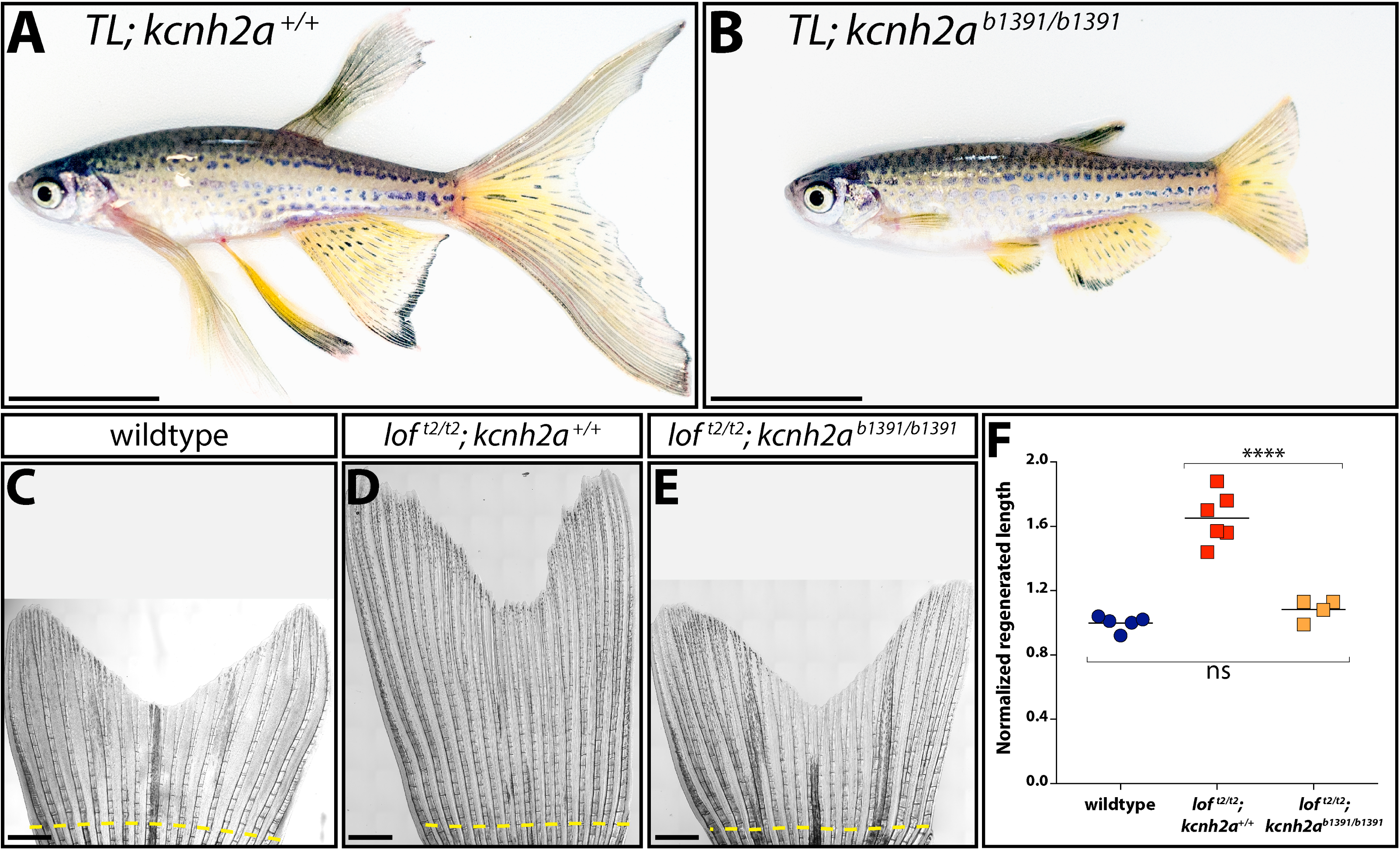
Fin overgrowth in *lof^t2^* requires *kcnh2a*. **(A, B)** Brightfield images of clutchmate *TL; kcnh2a^+/+^* (panel A) and *TL; kcnh2a^b1391/b1391^* (panel B) adult zebrafish. The scale bars represent 1 cm. **(C-E)** Representative images of regenerated caudal fins at 27 days post-amputation (dpa) from (C) wildtype control, (D) *lof^t2/t2^; kcnh2a^+/+^*, and (E) *lof^t2/t2^; kcnh2a^b1391/b1391^* fish. The dashed yellow line indicates the site of fin resection and scale bars represent 1 mm. **(F)** Quantification of fin ray outgrowth of wildtype (blue circles), *lof^t2/t2^; kcnh2^+/+^* (red squares), and *lof^t2/t2^; kcnh2a^b1391/b1391^* (orange squares) fish at 27 dpa. Each data point represents the normalized length of ray 3 from an individual animal of the indicated genotype. Asterisks indicate p < 0.005 determined by a one-way ANOVA with Tukey’s multiple comparisons tests. ns: not significant.

### Kcnh2a actively promotes fin overgrowth by slowing outgrowth termination

We measured regenerating fin outgrowth of wildtype and clutchmate *lof^t2^* fish over a 30 day period to explore temporal effects of ectopic *kcnh2a* on fin overgrowth. For each genotype, regenerated lengths over time fit well to a logistic growth curve, consistent with a lag for blastema establishment prior to a prolonged outgrowth phase. Wildtype and *lof^t2^* fish had indistinguishable maximal growth rates reached between 3 and 5 dpa (Figs. 3A, B and S1). Growth rates then gradually declined, approximating an exponential decay equation. However, the growth rate declined slower in *lof^t2^* fish, progressively increasing the fin length difference with wildtype animals. Wildtype fins largely ceased regenerative outgrowth by 21 dpa whereas *lof^t2^* fins continued to extend through the 30 dpa period. This prolonged outgrowth dynamic matches that of *lof^t2^* fin development (Iovine and Johnson, 2000). We conclude fin overgrowth in *lof^t2^* fish is due to an extended outgrowth period beginning around 6 dpa rather than elevated maximum growth.

**Figure 3.**
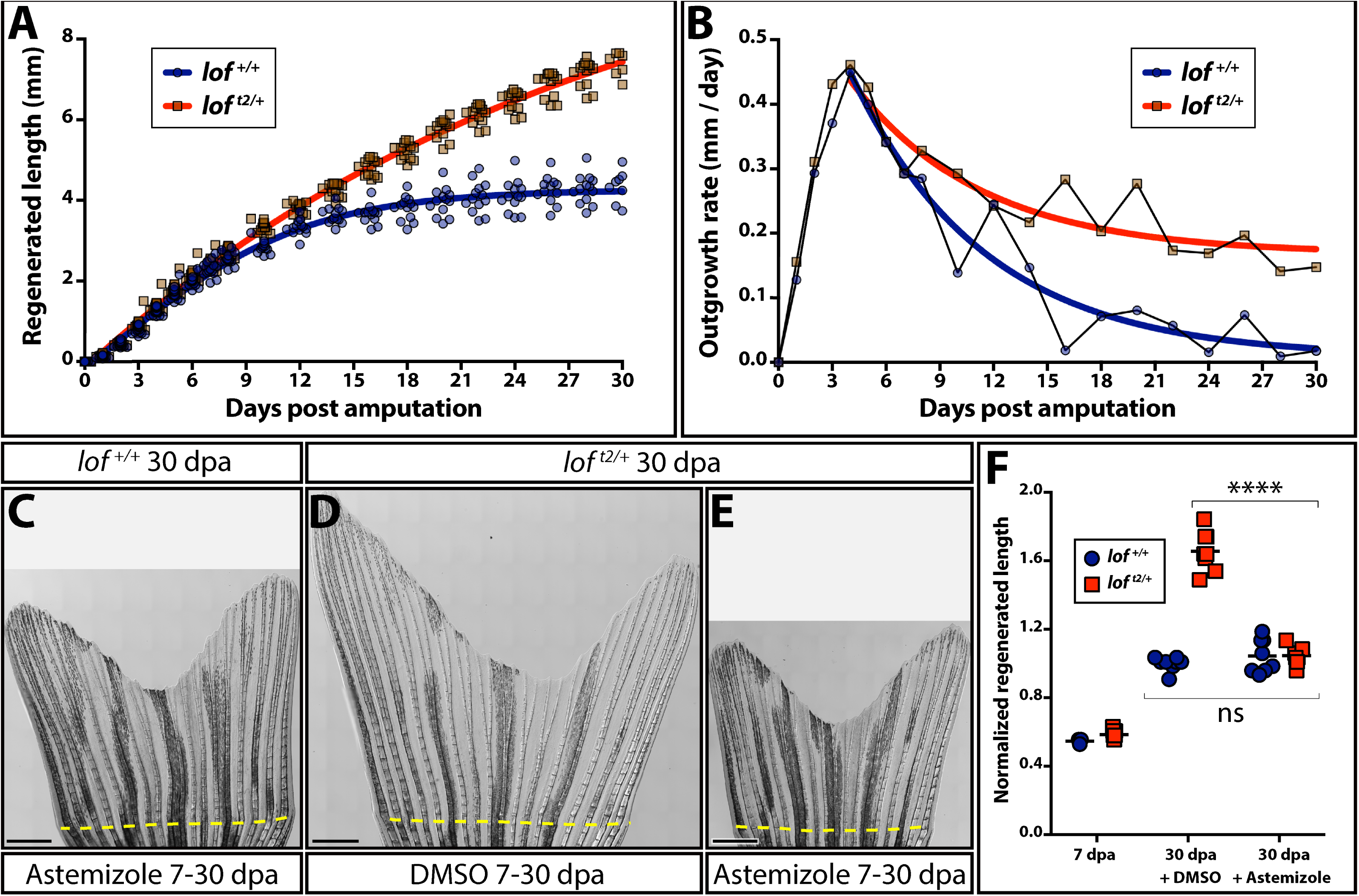
Kcnh2a actively prolongs the fin outgrowth period in *lof^t2^*. **(A)** Plots comparing the growth (A) and growth rate (B) of regenerating *lof^t2/+^* (blue) and clutchmate *lof^+/+^* (red) caudal fins. Curves show actual data fit to a (A) logistic equation reflecting establishment and then progressively slowing outgrowth phases and (B) exponential decay equation using time points after the peak growth rate is reached. All data points (≥ 12 fish per time) are shown in panel A and mean values are used in panel B. **(C-E)** Stitched DIC images showing clutchmate *lof^+/+^* astemizole-treated (panel C; 500 nM), DMSO-treated *lof^t2/+^* (panel D) and *lof^t2/+^*, astemizole-treated (panel E; 500 nM) fish at 30 days post caudal fin amputation (dpa). The dashed yellow line indicates the amputation plane and the scale bars denote 1 mm. **(F)** Quantification of normalized caudal fin regenerative outgrowth from control and astemizole-treated animals. Data points represent individual *lof^+/+^* (blue circles) and *lof^t2/+^* (red squares) animals prior to treatment (7 dpa) and post-treatment (30 dpa) with DMSO or astemizole (500 nM). Graph shows mean lengths of the third ventral regenerated ray normalized to DMSO-treated controls at 30 dpa. Each point is an individual animal. p < 0.0001 DMSO-treated from 7-30 dpa comparing wildtype and *lof^t2/+^* fin lengths, no significant differences between genotypes at 7 dpa or comparing 7-30 dpa astemizole-treated wildtype and *lof^t2/+^* regenerative outgrowth. Statistical tests used one-way ANOVA with Tukey’s multiple comparisons tests.

Kcnh2a is related to *ether-a-go-go* voltage-gated K^+^ channels (Vandenberg et al., 2012) whose channel activity is blocked by the small molecule astemizole (Sanguinetti and Tristani-Firouzi, 2006; Suessbrich et al., 1996; Zhou et al., 1999). We treated *lof^t2/+^* fish with astemizole from 7-30 dpa to determine if ectopic Kcnh2a actively extends fin outgrowth period. This inhibitor regimen fully suppressed excessive *lof^t2/+^* fin outgrowth to wildtype levels (Fig. 3C-F; p < 0.0001). Further, starting astemizole treatment at 6 dpa was as effective as administration throughout regeneration (Fig. S2). Therefore, although *lof^t2/+^* fins express *kcnh2a* relatively early during regeneration (4 dpa, Fig. 1), its functional impact on growth at these times is negligible. Further, consistent with wildtype-sized *kcnh2a^b1391/b1391^* caudal fins, Kcnh2a activity normally does not contribute to regenerative outgrowth. We conclude ectopic Kcnh2a exclusively causes regenerative fin overgrowth by actively preventing growth termination during late-stage outgrowth (∼7 dpa and onward).

### Ectopic *kcnh2a* acts in the intra-ray mesenchyme lineage to promote fin overgrowth in *lof^t2^*

Kcnh2a could function fin autonomously or systemically (e.g., via circulating endocrine factors) to prolong *lof^t2^* fin outgrowth periods. We used F_0_ CRISPR/Cas9 targeting to induce mosaic *kcnh2a* mutations to discriminate between these scenarios. The tissue autonomous model predicts fin size heterogeneity depending on the extent of *kcnh2a* targeting across rays of a given fin and between different fins. In contrast, the systemic hypothesis anticipates *kcnh2a* loss-of-function mutations always would equally suppress *lof^t2/+^* fin overgrowth. We injected embryos from a wildtype x *lof^t2/+^* cross with Cas9 and a *kcnh2a*-targeting gRNA and scored reared adults for fin overgrowth. Roughly half of uninjected control animals displayed long fins, as anticipated (51:40 wildtype:overgrown). In contrast, 21.8% of *kcnh2a* crispants showed partial, regionalized fin overgrowth (17 of 78, of which half would be *lof^t2/+^* fish). A representative partially overgrown caudal fin is shown in Fig. 4A. Another striking example displayed one pectoral fin rescued to wildtype length with the other exhibiting pronounced overgrowth indicative of *lof^t2^* (Fig. 4B). Therefore, ectopic Kcnh2a likely functions within fin tissue rather than systemically to disrupt the cessation of fin outgrowth.

**Figure 4.**
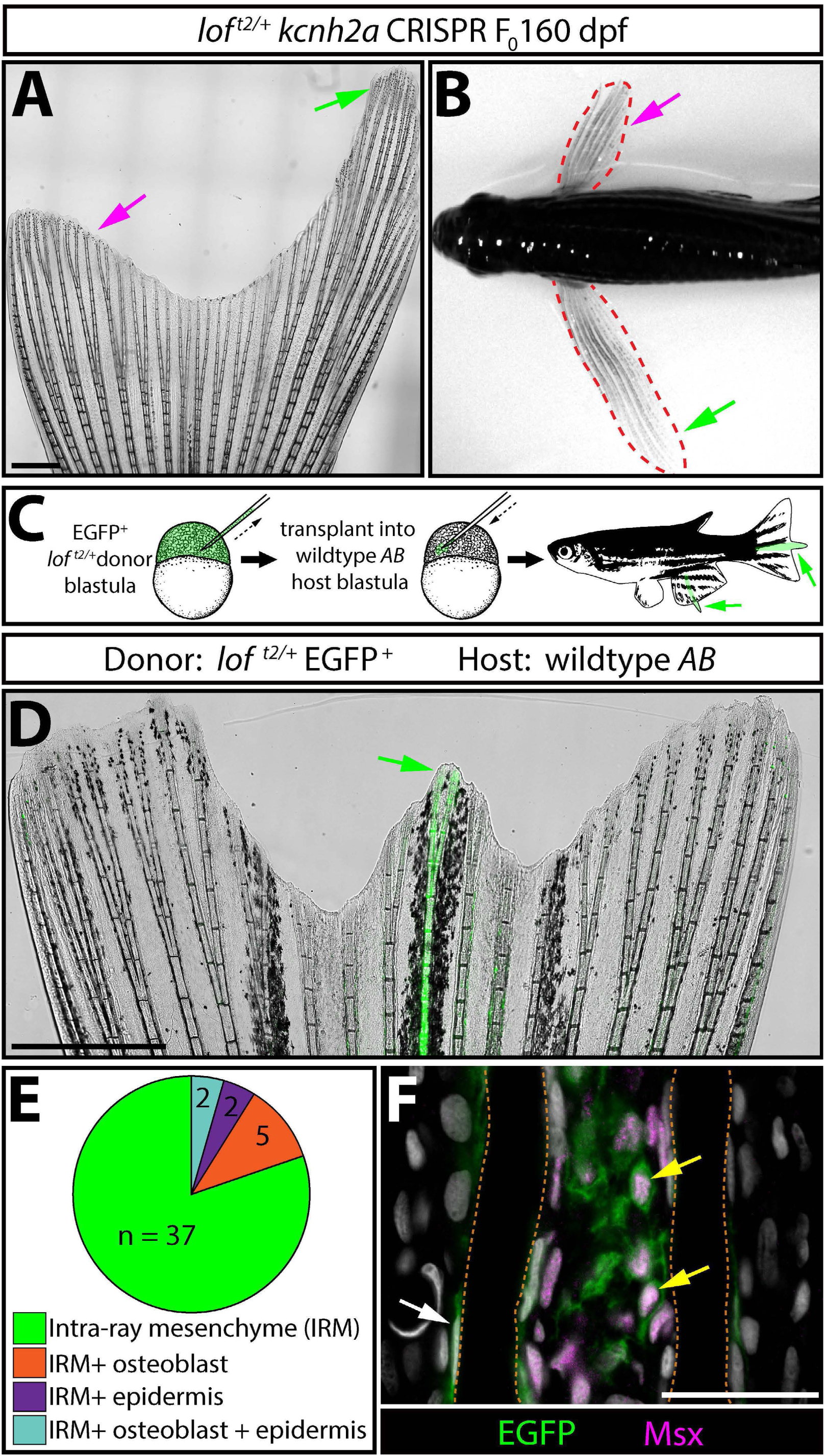
*kcnh2a* acts fin autonomously and within the intra-ray mesenchyme lineage to promote overgrowth. **(A)** Representative whole mount image of a F0 *kcnh2a* CRISPR *lof^t2/+^* adult caudal fin at 160 days post-fertilization. The magenta arrow points to suppressed overgrowth in otherwise long fins (green arrow). The scale bar represents 1 mm. **(B)** Pectoral fin asymmetry in a *kcnh2a* F_0_ CRISPR *lof^t2/+^* animal. The magenta arrow points to a wildtype-sized fin indicative of phenotypic suppression. The green arrow points to the overgrown contralateral fin expected of the *lof^t2/+^* genotype. **(C)** Upper panel schematic highlighting the nature of the blastula stage transplant experiment. EGFP^+^ *lof^t2/+^* blastula cells were transplanted into wildtype *AB* embryos. Reared adults with partial or complete fin overgrowth were scored for cell type(s) with EGFP expression. **(D)** A representative example of a caudal fin displaying overgrown EGFP^+^ fin rays. The green arrow points to overgrown EGFP^+^ fin rays and the scale bar represents 1 mm. **(E)** The pie chart indicates the EGFP^+^ lineage(s) present in 39 overgrown regions across 30 total fins with extended rays. **(F)** Caudal fin section of an overgrown chimeric fin ray immunostained with EGFP and Msx antibodies. Yellow arrows point to EGFP^+^/Msx^+^ intra-ray mesenchymal cells. The white arrow highlights an EGFP^+^ osteoblast. Fin rays are outlined with a dashed orange line. Hoechst-stained nuclei are in gray. The scale bar represents 50 µm.

We generated chimeras by blastula cell transplantations (Kimmel et al., 1990) to examine *lof^t2^* tissue autonomy further. We introduced EGFP-labeled *lof^t2/+^* blastula-stage cells into *AB* wildtype host embryos and raised them to adulthood (Fig. 4C). 13% of transplants (39 fins with overgrowth from 28 of 214 screened chimeras) exhibited EGFP^+^ fin tissue with notable overgrowth indicative of chimerism (Fig. 4D and Fig. S3). Several fins had strong EGFP^+^ overgrown rays flanked by intermediate length rays with minimal EGFP expression. We attribute these neighbor effects to diffusible factors emanating from the primary overgrown ray, anatomical influences on growth between adjacent rays, or scarce EGFP^+^ cells sufficient to cause lesser overgrowth. In a few cases, chimeric animals displayed asymmetrically sized pectoral fins, as observed with *kcnh2a* CRISPR-targeted F_0_ *lof^t2^* mosaic fish (Fig. S3). All overgrown chimeric rays contained EGFP^+^ intra-ray mesenchymal cells (n=30 overgrown fins containing only EGFP^+^ mesenchyme; Fig. 4E), although some also displayed EGFP-expressing osteoblasts and/or epidermis (n=5 EGFP^+^ mesenchyme/osteoblast; n=2 EGFP^+^ mesenchyme/epidermis; n=2 EGFP^+^ mesenchyme/osteoblast/epidermis Fig. 4E). We immunostained sections from representative overgrown fins for the mesenchymal marker Msx (Akimenko et al., 1995) to confirm overgrown chimeric fins always included EGFP^+^ *lof^t2/+^*-derived intra-ray mesenchyme (Fig. 4F and Fig. S3). In contrast, chimeras harboring only EGFP^+^ *lof^t2/+^* epidermal cells were of normal length (n=3, Fig. S4). We conclude ectopic Kcnh2a functions in intra-ray mesenchyme and/or blastema cells derived from this lineage to disrupt outgrowth-slowing mechanisms.

We used RNAscope (Wang et al., 2012) to localize *kcnh2a* mRNA in 4 dpa *TL* caudal fin sections. As predicted from our transplant studies, *kcnh2a* was expressed in *TL* regenerating intra-ray mesenchyme, including at lower levels in growth-promoting distal blastema cells of shared lineage (Tornini et al., 2017; Wehner et al., 2014) (Fig. 5A). We did not detect *kcnh2a* in *AB* control fins, as expected from RNA-Seq and qRT-PCR (Fig. 5B). We next combined *kcnh2a* RNAscope with immunostaining for the intra-ray mesenchyme lineage markers Msx (Akimenko et al., 1995) or *tph1b:mCherry* (Tornini et al., 2016). *kcnh2a* was expressed in Msx^+^ or *tph1b:mCherry^+^* cells in proximal regenerating tissue (Fig. 5C-F) with lower levels in *tph1b:mCherry^+^* distal blastema cells (Fig. S5), reinforcing that ectopic Kcnh2a acts in this lineage to promote fin overgrowth.

**Figure 5.**
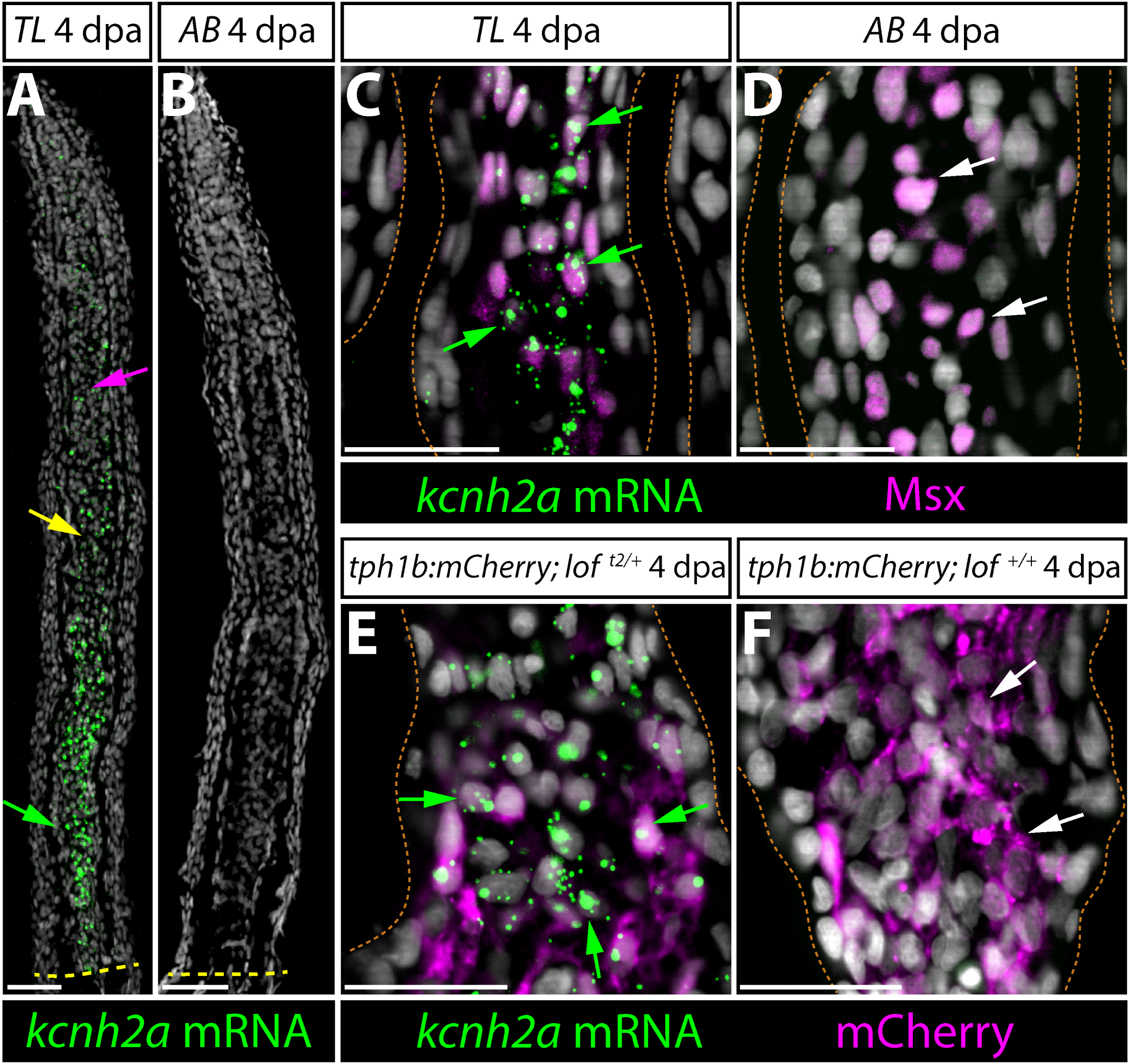
*kcnh2a* is expressed ectopically in *lof^t2^* intra-ray mesenchyme lineage cells during fin regeneration. **(A, B)** *kcnh2a* mRNA localization (in green) detected by RNAscope in longitudinal caudal fin sections from 4 day post amputation (dpa) *TL* (*lof^t2/t2^*, panel A) and *AB* (panel B) animals. The green arrow points to proximal intra-ray cells expressing high levels of *kcnh2a*, which is also detected in medial mesenchyme (yellow arrow) and distal cells (magenta arrow) of *TL* fish. The dashed yellow line denotes the site of amputation. Hoechst stained nuclei are in gray and scale bars indicate 50 µm. **(C, D)** Double *kcnh2a* RNAscope (green) and Msx immunostaining (magenta) of 4 dpa caudal fin sections from *TL* (panel C) and *AB* (panel D) fish. **(E, F)** Combination *kcnh2a* RNAscope (green) and mCherry immunostaining (magenta) of 4 dpa fin sections from (E) *tph1b:mCherry; lof^t2/+^* and (F) *tph1b:mCherry; lof^+/+^* fish. For panels C-F, green arrows highlight Msx^+^ or *tph1b:mCherry*^+^ cells with overlapping *kcnh2a* mRNA in proximal regenerating *lof* tissue. White arrows show Msx^+^ or *tph1b:mCherry*^+^ nuclei in corresponding regions from control fins lacking *kcnh2a* expression. Fin rays are outlined with a dashed orange line. Hoechst-stained nuclei are in gray. Scale bars represent 50 µm.

Intra-ray mesenchymal cell proliferation rates monitored by EdU incorporation were similar in 4 dpa wildtype and *lof^t2/+^* regenerating fins (Fig. S6). Cell cycling was distally concentrated in both genotypes. The fraction of proliferating cells was significantly higher in 10 dpa *lof^t2/+^* fin regenerates, being only modestly reduced from 4 dpa (Fig. S6). EdU-incorporating intra-ray cells remained largely confined to the distal blastema of *lof^t2/+^* fin regenerates in spite of ectopic *kcnh2a* throughout ray mesenchyme. Further, the *wnt5a* growth factor remained distal-enriched in 10 dpa *lof^t2/+^* regenerating fins (Stoick-Cooper et al., 2007, Stewart et al., 2014; Fig. S6). Therefore, matching our outgrowth rate measurements, an extended outgrowth period leading to accumulated cell proliferation rather than an increased maximum cycling rate causes *lof^t2^* fin overgrowth. Further, *kcnh2a* prolongs outgrowth without being autonomously sufficient to drive cell cycling and/or a growth factor-producing distal blastema cell state.

### Calcineurin promotes outgrowth cessation to help establish fin size

Inhibiting the Ca^2+^/calmodulin-dependent phosphatase calcineurin with the immunosuppressants FK506 or cyclosporin leads to pronounced fin overgrowth (Daane et al., 2018; Kujawski et al., 2014). Maximal growth rates in calcineurin-inhibited regenerating caudal fins are the same as wildtype (Daane et al., 2018; Kujawski et al., 2014), implying that, like *lof^t2/+^*, such fins do not properly terminate growth. We treated fish with FK506 at various time points post-caudal fin amputation to test this hypothesis. Optimization experiments demonstrated that daily 4 hour immersion from 1-21 dpa in water containing 500 nM FK506 was sufficient to overgrow fins to the same degree as *lof^t2/t2^* with no apparent adverse effects on animal health (Fig. S7). We treated regenerating animals from either 1-5, 6-21, or 1-21 dpa following this acute drug delivery regimen and measured regeneration extents at 21 dpa. Treating animals with FK506 from 1-5 dpa had no effect on fin outgrowth but still prevented joint formation (Fig. 6A, B), a known additional phenotype upon calcineurin inhibition (Kujawski et al., 2014). In contrast, FK506 treatment from 6-21 or 1-21 dpa caused the same extent of pronounced fin overgrowth (Fig. 6C, E). Thus, like ectopic Kcnh2a in *lof^t2^* mutants, calcineurin inhibition appears uniquely to cause fin overgrowth by disrupting growth cessation.

**Figure 6.**
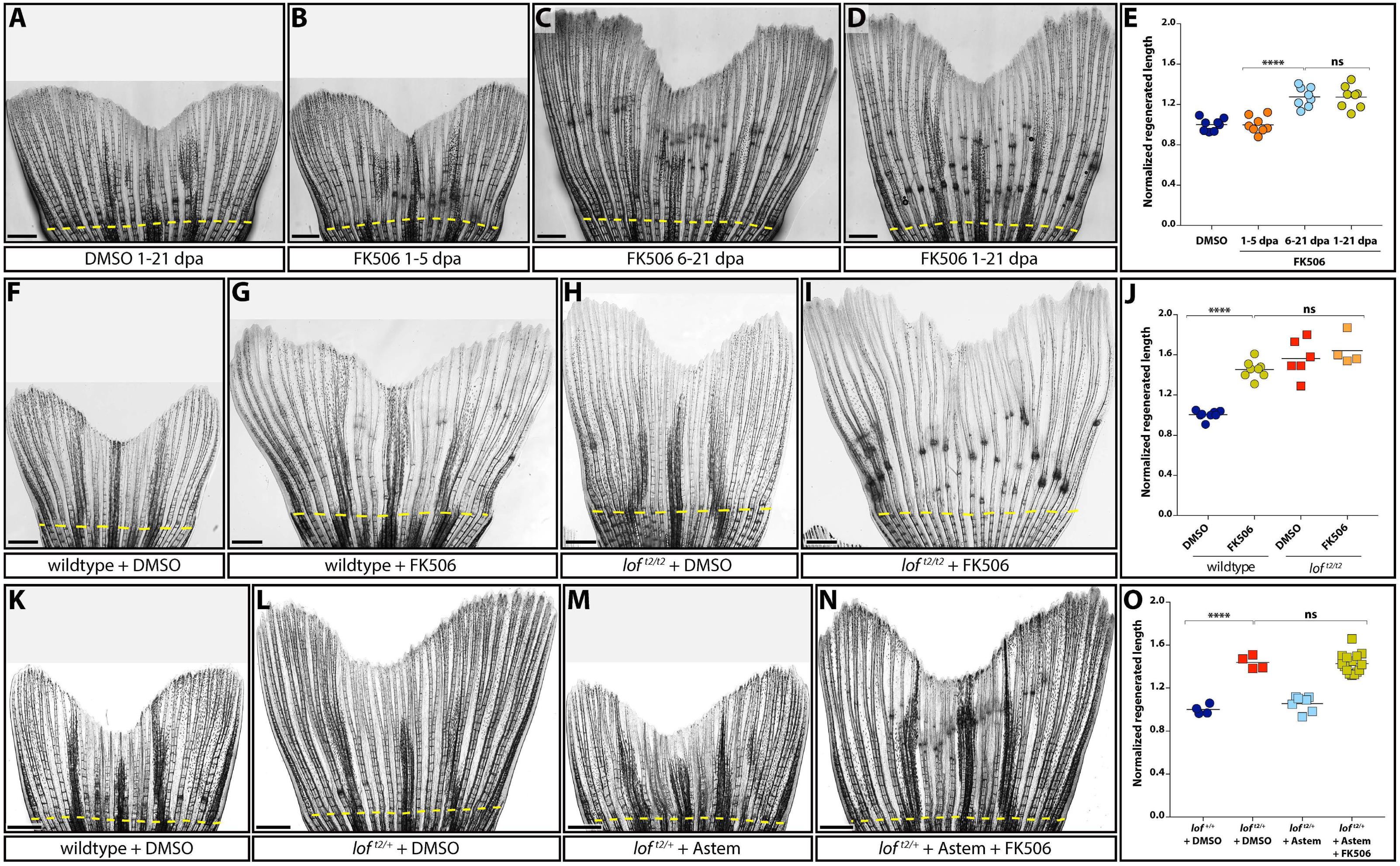
Kcnh2a-disrupted calcineurin signaling gradually terminates fin outgrowth. (**A-D**) Calcineurin gradually ends the fin outgrowth period. Stitched DIC images showing wildtype 21 day post amputation (dpa) caudal fins treated with DMSO (A) or 500 nM FK506 from either 1-5 (B), 6-21 (C), or 1-21 dpa (D). **(E)** Quantification of the experiment presented in A-D. Data are the average regenerated lengths of the third ray normalized to DMSO-treated controls at 21 dpa. Each point is a single wildtype animal treated with either DMSO 1-21 dpa (dark blue circles) or FK506 for the indicated times (orange circles 1-5 dpa, light blue circles 6-21 dpa, gold circles 1-21 dpa). **(F-I)** Ectopic Kcnh2a and FK506 treatment do not cooperate during overgrowth. DIC-imaged regenerated caudal fins from wildtype (F, G) and *lof^t2/t2^* (H, I) animals treated with DMSO (F, H) or 500 nM FK506 (G, I) daily from 1-23 dpa. **(J)** Graph showing relative regenerate lengths of ray 3 at 23 dpa from wildtype fish treated with DMSO or FK506 (blue or gold circles, respectively), and *lof^t2/t2^* fish treated with DMSO or FK506 (red or orange squares respectively). Each data point is an individual fish. **(K-N)** Kcnh2a activity is not required for FK506-induced overgrowth. Images showing wildtype (K) and *lof^t2/+^* clutchmate (L-N) caudal fin regenerates at 21 dpa. Animals were treated from 7-21 dpa with DMSO vehicle (K, L), 500 nM astemizole (M), or 500 nM astemizole + 500 nM FK506 (N). **(O)** Plot showing relative regenerative fin growth of ray 3 from the samples described in (K-N). Data points represent individual animals: DMSO-treated *lof^+/+^* or *lof^t2/+^* (dark blue circles and red squares, respectively), astemizole-treated *lof^t2/+^* (light blue squares), and *lof^t2/+^* treated with both FK506 and astemizole (gold squares). Data are normalized to DMSO-treated wildtype samples. Dashed yellow lines indicate amputation sites. Each data point represents an individual animal and asterisks indicate p < 0.001; ns: not significant. Scale bars are 1 mm. One-way ANOVA with Tukey’s multiple comparisons tests tested differences between regenerated fin lengths.

### Kcnh2a in *lof^t2^* likely disrupts calcineurin signaling to cause fin overgrowth

We hypothesized ectopic Kcnh2a disrupts calcineurin signaling given *lof^t2^* and FK506 experiments highlight similar growth dynamic effects and temporal functions during fin regeneration. If so, *lof^t2^* should not enhance FK506-induced fin regenerative overgrowth. Accordingly, we observed no difference in caudal fin overgrowth in wildtype vs *lof^t2/t2^* fish treated with FK506 from 7-21 dpa (Fig. 6F-J). The *lof^t2/t2^* genetic background did not alter FK506 bio-availability because joint formation remained dramatically disrupted (Fig. 6H, I). Additionally, treating *lof^t2/+^* with both astemizole and FK506 from 7-21 dpa still produced FK506-induced fin overgrowth (Fig. 6K-O; p <0.001). Therefore, Kcnh2a activity, unlike Kcnk5b/*alf* (Daane et al., 2018), is not downstream of calcineurin. Further, *lof^t2^*-ectopic Kcnh2a throughout fin development, and even the first week of fin regeneration, does not impact how an amputated fin responds to calcineurin inhibition. We conclude ectopic Kcnh2a likely disrupts calcineurin signaling, which otherwise impacts fin size by gradually terminating outgrowth.

## DISCUSSION

### *lof^t2^* is a neomorph allele of *ether-a-go-go* related *kcnh2a* potassium channel

*lof^t2^* was one of the first zebrafish mutant lines used scientifically (hence, its “t2” – Tübingen 2 – designation), having been originally isolated by tropical fish hobbyists (Elias, 1984; Haffter et al., 1996; Van Eeden et al., 1996). *lof^t2^* remains widely used because its highly specific fin overgrowth provides a convenient phenotypic marker to discriminate zebrafish of mixed genotypes. We demonstrate *lof^t2^* is a regulatory neomorphic allele of *kcnh2a* (*kcnh2a^lof^*) that causes its ectopic expression in fin mesenchyme, resolving the basis of its remarkable and long-appreciated phenotype. The genetic lesion is likely a chromosomal inversion linking *kcnh2a* to a displaced enhancer, explaining why *kcnh2a* lies just outside of the originally mapped *lof* region (Daane et al., 2021; Iovine and Johnson, 2000).

*kcnh2a* encodes an *ether-a-go-go* (EAG)-related voltage-gated K^+^ channel. EAG/KCNH channels support membrane repolarization in various excitable cells, including neurons and myocytes (Vandenberg et al., 2012). The human ortholog KCNH2 produces *I_Kr_*, the rapid component of the cardiac delayed rectifier current (Curran et al., 1995; Noble and Tsien, 1969; Sanguinetti et al., 1995). The voltage-dependent gating properties of KCNH2, namely slow opening and closing but fast inactivation, uniquely enables it to repolarize cardiac tissue and terminate action potentials (Bohnen et al., 2017; Vandenberg et al., 2012). *KCNH2* mutations are a frequent cause of inherited arrhythmias known as long QT syndrome whereby patients exhibit prolonged cardiac action potentials (Bohnen et al., 2017; Curran et al., 1995; Sanguinetti et al., 1995). KCNH2 is also a notorious pharmaceutical “off-target”, leading to withdrawal of many drugs due to arrhythmia side effects. We find homozygous loss of *kcnh2a* has no overt effects on zebrafish development to adulthood, including fin length, although we have not assessed cardiac function. Ectopic expression of *kcnh2a* in *lof^t2^* may be confined to fin mesenchyme, explaining the highly specific phenotype and otherwise healthy fish. However, we have not explored if *kcnh2a* is misexpressed in other *lof^t2^* tissues, where it could adversely impact cardiac or other excitable cell functions. If so, *lof^t2^* likely should be avoided as a “wildtype” strain.

### Size establishment by modulation of outgrowth periods by intra-ray mesenchymal cells

Our study of fin overgrowth in *lof^t2^* advances a mechanistic view of fin growth control centered on growth cessation rather than acceleration or potency. Ectopic Kcnh2a in *lof^t2^* does not alter initial or maximal growth rates during caudal fin regeneration, similar to *lof^t2^* fin development growth dynamics (Iovine and Johnson, 2000). Rather, Kcnh2a expression dampens growth rate deceleration, effectively extending the allometric growth period. Kcnh2a specifically may enhance later outgrowth phases because mitogenic drive is saturated during the initial regenerative growth response (3-6 dpa), overshadowing Kcnh2a growth-promoting effects. Alternatively, upstream signals activating Kcnh2a or the ion signaling pathways disrupted by Kcnh2a may be inactive early in regeneration.

Our CRISPR and transplant chimera studies indicate ectopic Kcnh2a within fins and their intra-ray mesenchyme lineage is necessary and sufficient for fin overgrowth. Concordantly, *lof^t2^* fish ectopically express *kcnh2a* in blastema fin mesenchyme but not other cell types with transcripts nearly undetectable in wildtype regenerating caudal fins. *alf* transplant experiments suggest hypermorphic Kcnk5b also acts within the mesenchyme lineage to cause fin overgrowth (Perathoner et al., 2014). Further evidence indicating this population is growth-determining include lineage tracing experiments showing intra-ray mesenchyme contributes to the growth-promoting distal blastema (Tornini et al., 2016). Finally, we recently proposed a model explaining slowing fin outgrowth by the progressive depletion of these distal blastema cells by biased differentiation vs. self-renewal (Stewart et al., 2019). Therefore, Kcnh2a may disrupt ion signaling that normally promotes mesenchyme lineage cell transitions from a growth-promoting to differentiated state.

Alternatively, ectopic Kcnh2a may prolong or enhance the production of pro-growth signaling molecules during the slowing outgrowth phase of fin regeneration. In support, Wnts and FGFs are produced by mesenchyme-derived “organizing center” or “niche” cells at the distal blastema (Stewart et al., 2019; Tornini et al., 2016; Wehner et al., 2014) and new observations suggest the Kcn5b (*alf*) K^+^ channel cell autonomously promotes growth factor production (Yi et al., 2020). We observed *kcnh2a* expression throughout regenerating mesenchyme, highest near the amputation site but still detectable in medial and distal blastema. However, we did not observe elevated proliferation or ectopic *wnt5a*, a representative distal growth factor, in proximal *kcnh2a*-expressing *lof^t2^* mesenchyme. Nevertheless, ectopic Kcnh2a could prolong growth factor production by acting directly in distal blastema/niche cells or in more proximal mesenchyme by disrupting negative feedback to the distal cells.

### Calcineurin as an ion signaling node for growth cessation

Mutations in *kcnk5b* (K^+^ channel, (Perathoner et al., 2014)), *kcc4a*/*slc12a7A* (K^+^ Cl^-^ co-transporter, (Lanni et al., 2019)), or over/ectopic expression of the K^+^ channels *kcnj13, kcnj1b*, *kcnj10a*, *kcnk9c* (Silic et al., 2020), and now *kcnh2a* (this study) all cause fin overgrowth. Each model may disrupt a common “ion signaling” pathway featuring the fin outgrowth-restraining Ca^2+^-dependent phosphatase calcineurin (Daane et al., 2018; Harris et al., 2020; Kujawski et al., 2014; Lanni et al., 2019; Yi et al., 2020). Here, we found fin overgrowth in regenerating *lof^t2^* fins was not enhanced by the calcineurin inhibitor FK506, suggesting ectopic Kcnh2a also inhibits calcineurin signaling. Ectopic Kcnh2a in *lof^t2^* could derail calcineurin output either upstream or downstream of calcineurin itself. Supporting the former, calcineurin’s phosphatase activity is modulated by sustained elevated cytosolic Ca^2+^ (Klee et al., 1998; Rao, 2009; Timmerman et al., 1996) and KCNH2 effectively shortens Ca^2+^ fluxes during the cardiac conduction cycle (Bohnen et al., 2017; Vandenberg et al., 2012). Further, recent work indicates reduced calcineurin activity in *lof^t2^* fins (Cao et al., 2021). Alternatively, ectopic Kcnh2a could short-circuit calcineurin-promoted ion signaling dynamics mediated by Kcnk5b inhibition (Daane et al., 2018; Yi et al., 2020). These possibilities could be distinguished by determining if expressing constitutively active calcineurin in the intra-ray mesenchyme lineage suppresses *lof^t2^* fin overgrowth.

### Ion signaling and interpretation of size-instructing positional information

Bioelectricity is widely linked to organ size control and regeneration (McLaughlin and Levin, 2018). Bioelectric fields are suggested to pre-pattern undifferentiated tissue (including the fin blastema) to establish positional information instructing correct amount of growth. However, *lof^t2^* does not seem to change positional information established at the outset of fin regeneration because inhibiting ectopic Kcnh2a only during the late outgrowth phase restores a normal sized fin. Likewise, calcineurin need only be inhibited late during regeneration to maximally overgrow fins. Therefore, elevated Kcnh2a and calcineurin inhibition appear to disrupt growth deceleration mechanisms tuned to interpret, rather than set, positional information and thereby help establish (and re-establish) correct proportions. Regeneration has the additional challenge that outgrowth has to “read” some form of memory within cells or in higher tissue-level organization to direct correct amount of outgrowth while also re-establishing said memories. At least proximally, *lof^t2^* overgrown caudal fins do not carry abnormal positional memory as fins amputated here regenerate normally when ectopic Kcnh2a is inhibited. Similarly, clonal analyses show calcineurin inhibition does not alter blastema pre-patterning (Tornini et al., 2016) and re-amputation of previously FK506-treated animals results in normal fin size (Daane et al., 2018). The nature of fin positional information and memory is unresolved but, as mentioned, likely reflects properties of intra-ray fibroblasts and derived blastema mesenchyme cells and/or their population sizes (Stewart et al., 2019).

Identifying and characterizing ectopic *kcnh2a* as the cause of the classic *lof^t2^* zebrafish allele provides a framework to consider bioelectric control of organ size and shape through ion-mediated intracellular signaling. Subtle changes in calcium dynamics that tune signaling output and then growth period durations could produce profound changes in organ scale while retaining overall form and function. More local expression changes in ion signaling components (therefore, acting as effectors of “positional information”) could then readily alter organ proportions supporting evolutionary innovations and phenotypic diversity. Extending this concept, intersecting systemic gene regulatory signals including hormones could underlie sexual dimorphic traits or environmental effects on organ morphology arising after embryonic development. Swordtail fish provide a compelling example by their male-specific, dramatically elongated rays (the “sword”) on the ventral edge of the caudal fin. Strikingly, the swordtail phenotype was recently linked to the *kcnh2a*-related gene *kcnh8* (Schartl et al., 2020). How fin growth periods are highly sensitive to alterations in K^+^ channels and Ca^2+^/calcineurin is unclear. Key next steps include characterizing membrane potentials, putative depolarizing signals, and how Ca^2+^ signaling alters states and/or activities of intra-ray mesenchymal cells during fin outgrowth.

### MATERIALS AND METHODS

#### Zebrafish strains and maintenance

Zebrafish were housed in the University of Oregon Aquatic Animal Care Services facility at 28-29°C. The University of Oregon Institutional Animal Care and Use Committee oversaw animal use. Wildtype *AB* (University of Oregon Aquatic Animal Care Services), *TL* (Haffter et al., 1996), *Tg(tph1b:mCherry)^ens700Tg^* (Tornini et al., 2016), *Tg(sp7:EGFP)^b1212Tg^* (DeLaurier et al., 2010), and *Tg(Xla.Eef1a1-actb2:LOXP-LOX5171-FRT-F3-EGFP,mCherry)^vu295aTg^* (Boniface et al., 2009), abbreviated herein *Tg(eab:FlEx),* zebrafish lines were used.

#### CRISPR/Cas9-generation of *kcnh2a* mutants

A guide RNA (gRNA) encompassing the putative *kcnh2a* start codon (in bold) 5’ GACAAC**ATG**CCTGTACGACG 3’ was synthesized in vitro and co-injected with recombinant Cas9 protein (500 ng/µl; Thermo Fisher) into one-cell stage *TL* embryos. F0 fish were outcrossed and those F1 clutches harboring mutant alleles identified by PCR sequencing reared to adulthood. Genotyping of F1 adults identified a predicted nonsense allele, *kcnh2a^b1391^*, that deletes 8 bp (underlined) at the gRNA target site (GACAAC**ATG**CCT<u>GTACGACG</u>) to produce a frameshift at codon 3 of the predicted polypeptide and remove a Hpy99I restriction endonuclease site. *kcnh2a^b1391^* was genotyped by amplifying genomic DNA using primers *kcnh2a*_P1 and *kcnh2a*_P2 and digesting PCR products with Hpy99I.

To distinguish between a tissue autonomous vs. systemic function for *kcnh2a* in *lof^t2/+^* fin outgrowth, CRISPR/Cas9 mutagenesis was carried out on one-cell stage embryos from a *lof^t2/+^* x *lof^+/+^* cross. These animals were raised to adulthood and examined for fin ray length heterogeneity within fins and fin size disparities between paired fins.

#### Sequencing of *kcnh2a* in *longfin^t2^*

*kcnh2a* coding exons were PCR amplified from *TL* genomic DNA (primers listed below), ligated to a vector (pCRII, Thermo Fisher), and sequenced using T7 and SP6 primers. Additional sequencing primers were used to ensure complete coverage of the clone.

#### Adult whole zebrafish and fin imaging

Whole animal images were captured using a homemade light box made from a fenestrated styrofoam container, a halogen bicycle lamp, and a consumer Fujifilm X-A1 camera with Fujinon 28mm 1.4R lens. High resolution fin images were obtained from tricaine-euthanized adult fish placed in water or mounted in 0.75% low melting agarose. Stitched differential interference contrast (DIC) images were then captured using a motorized Nikon Eclipse Ti widefield microscope with a 10x objective and NIS-Elements software.

#### RNA-Seq

Caudal fins of adult *lof^t2/+^* and *lof^+/+^ Tg(sp7:EGFP)* clutchmates were amputated. At 4 dpa, regenerated tissue beyond the GFP-marked differentiated osteoblast domain was homogenized in TRIzol reagent (Thermo Fisher). Tissue pooled from four animals constituted matched replicate samples. RNA was isolated following the manufacturer’s instructions with minor alterations. The RNA was precipitated overnight at −80°C and then pelleted at 15,000 rpm for 30 minutes at 4°C. Pellets were washed twice with 70% ethanol, dried for 10 minutes at room temperature, and resuspended in RNase-free water (Thermo Fisher). RNA-Seq libraries were prepared from 1 µg of isolated RNA using a Kapa Biosystems Stranded mRNA-Seq kit. Bar-coded libraries were pooled and sequenced using a HiSeq 4000 (Illumina) and reads were aligned to the zebrafish genome (GRCz11) using TopHat2 (Kim et al., 2013). Aligned reads were scored using HTseq (Anders et al., 2015) with edgeR (Robinson et al., 2010) used to identify differentially expressed transcripts.

#### RT-qPCR

RNA was isolated from whole caudal fin regenerates of adult *lof^t2/+^* and *lof^+/+^* clutchmates at 96 hpa. Each replicate represented a pooled sample of three fin regenerates. RNA was isolated as described in the preceding section. cDNA was synthesized from 500 ng of total RNA using oligo(dT)18 primer and Maxima H Minus RT (Thermo Fisher) following the manufacturer’s protocol. Primers *kcnh2a*_P1 and *kcnh2a*_P2 were used to determine *kcnh2a* expression. Primers for *runx2b* and *rpl8* were used as a control and ubiquitously expressed reference gene, respectively.

The *kcnh2a^AB^* polymorphism was identified serendipitously in University of Oregon Aquatic Animal Care Services facility stock AB fish and has been observed previously in other zebrafish strains (Butler et al., 2015). To demonstrate monoallelic expression of *kcnh2a* on mutant *lof^t2^* chromosome 2, we generated *lof^t2^ kcnh2a^WIK-TL^; lof^+^ kcnh2a^AB^* fish and prepared cDNA from 4 dpa caudal fins as described above. Quantitative PCR used the primer combinations: *kcnh2a*_P1 and *kcnh2a*_P2, *kcnh2a*_P1 and *kcnh2a^WIK-TL^*, and *kcnh2a*_P1 and *kcnh2a^AB^*. For all studies, qPCR was quantified by the ΔΔC_t_ method using *rpl13* mRNA levels for normalization as described (Stewart et al., 2014). RT-qPCR statistical analyses used a repeated measures one-way ANOVA with Tukey’s multiple comparisons test.

#### Small molecule treatments

For astemizole treatments, fins of *lof^t2/+^* and clutchmate animals were amputated and allowed to regenerate for the indicated times. When appropriate, astemizole (5 mM stock solution in DMSO) was added directly to fish water to a final concentration of 500 nM; control fish received an equal volume of DMSO. Fish were fed and changed to fresh drug- or DMSO-containing water every 48 hours. The unusually long half-life of astemizole (> 7 days in mammals, (Paton and Webster, 1985) precluded testing how exclusively early treatments affect outgrowth. For FK506 experiments, fins were amputated and animals treated with indicated concentrations of FK506 (typically, 500 nM; stock solution in DMSO) in static water for 4 hours every day for the indicated periods and then returned to normal water flow conditions. For Kcnh2a/calcineurin epistasis experiments, we combined treatments with astemizole and FK506 from 7 – 21 dpa for 4 hours in static water containing 500 nM astemizole +/-500 nM FK506, then returned to system water. At the experiment’s conclusion, each caudal fin was imaged on a Leica M205 FA stereomicroscope and lengths of the third ray from the amputation site to the fin’s tip were measured (detailed below). For each experimental group, representative animals were imaged by DIC stitched imaging as described previously.

#### Fin measurements

To quantify fin growth after regeneration, the length of the third ray was measured from the amputation site to the tip of the fin using stereomicroscope images and FIJI software (NIH). Growth measurements from individuals comprising each cohort were normalized to the mean of the control group. A one-way ANOVA with Tukey’s multiple comparisons tests tested differences between regenerated fin lengths. Logistic equation curves were fit to the fin regeneration length over time data to account for a slowly outgrowing establishment phase followed by rapid acceleration and then slow deceleration during the outgrowth phase. An exponential decay curve was fit to the outgrowth rate over time data starting from the peak growth rate at 4 dpa. Prism 7 was used for statistical analyses, curve fitting, and graph plotting (GraphPad Software).

#### Immunostaining and EdU staining

Fin tissue was paraffin embedded and sectioned as described (Stewart et al., 2014). Slides were antigen retrieved for 5 minutes in 1 mM EDTA, 0.1% Tween-20 using a pressure cooker followed by blocking in 10% milk in phosphate buffered saline containing 0.1% Tween-20. Antibodies were applied overnight in blocking buffer at 4°C. Individual antigens were visualized using Alexa dye-conjugated secondary antibodies (Thermo Fisher). Stained sections were imaged using Olympus FV1000 or Zeiss LSM880 laser scanning confocal microscopes.

For 5-ethynyl-2’-deoxyuridine (EdU) incorporation studies, caudal fins from *tph1b:mCherry* and clutchmate *tph1b:mCherry; lof^t2/+^* fish were collected at 4 or 10 dpa following a 4 hour pulse with 1 mg/kg EdU (Thermo Fisher) delivered by intra-peritoneal injection. Fins were processed for immunostaining as described above. Only the ventral lobe was collected for long finned fish. EdU detection was carried out according to the manufacturer’s recommendations (Thermo Fisher). Slides were imaged on a Zeiss LSM880 confocal microscope. EdU^+^ mesenchymal cells were quantified using the Imaris software package Spots function on a masked channel excluding non-blastema cells. The statistical analysis used a two-way ANOVA with Tukey’s multiple comparison tests.

#### RNAscope combined with EdU or immunostaining

RNAscope probes to detect *kcnh2a* or *wnt5a* mRNA were designed and synthesized by ACD Bio. RNAscope was performed using the Multiplex Fluorescent kit (ACD Bio) according to manufacturer’s recommendations for paraffin embedded sections with minor modifications. RNAscope followed by EdU was performed for dual imaging. Combined immunostaining using Msx (Developmental Studies Hybridoma Bank; 1:20 dilution of hybridoma supernatant) or mCherry (Sicgen; 1:100) antibodies was carried out after RNAscope as directed by the manufacturer (ACD Bio). Nuclei were visualized by Hoechst staining (Thermo Fisher). Imaging used Olympus FV1000 or Zeiss LSM880 laser scanning confocal microscopes.

#### Generation of chimeric animals

Blastula stage cell transplantations were performed as described (Kimmel et al., 1990). Briefly, 20-50 mesoderm-targeted single cells from high-stage *lof^t2/+^; Tg(eab:FlEx)* donors were transplanted into *AB* host high-stage embryos (Kimmel et al., 1995). Chimeric animals were reared to adulthood and fins screened for EGFP expression and overgrowth. Those with apparent fin overgrowth were imaged on a Leica M205 FA stereomicroscope. Stitched high-resolution images of select overgrown fins were also collected. Epifluorescent images of overgrown fins were examined to determine lineages including EGFP^+^ cells, as described (Stewart and Stankunas, 2012). Lineage-labeling of select samples was confirmed by immunostaining fin sections using EGFP (Aves Labs Inc.; 1:1000), Msx (Developmental Studies Hybridoma Bank; 1:20), sp7 (A-13; Santa Cruz Biotechnology; 20 ng/ml) and p63 (PA5-36069; Thermo Fisher; 1:100) antibodies as described earlier.

#### Primers

The following primers were used:

*kcnh2a*_P1 5’ GGATTTGCGCCCTCCGAACACTAAC 3’

*kcnh2a*_P2 5’ GTGCGCGCATCCTCGAGCTCAG 3’

*kcnh2a*_P3 5’ ACTGGAGACTTTGAAGAGACGC 3’

*kcnh2a*_P4 5’ GAATGATGGTGTCCAGAAAGGT 3’

*kcnh2a*^AB^ 5’ GTACAGGCATGTTTGTCCC 3’

*kcnh2a*^WIK-TL^ 5’ GTACAGGCATGTTGTCCT 3’

*kcnh2a*_exon_1-2_for 5’ CCCCAACCATCCTCTCACTGCCTCTCC 3’

*kcnh2a*_exon_1-2_rev 5’ CAGGGCCCCTAAGCTGTCCTGG 3’

*kcnh2a*_exon_3_for 5’ CTGGTGATCGCAGACCTTAACACAGACCTC 3’

*kcnh2a*_exon_3_rev 5’ GAATTGACCCCCTCCCTGATCGTCAACG 3’

*kcnh2a*_exon_4_for 5’ CTGGTGAACTGCTGCAAGTAAGTTACTGTAGATTC 3’

*kcnh2a*_exon_4_rev 5’ CTGTATGACCACTGTATGACCAGGTGTACC 3’

*kcnh2a*_exon_5_for 5’ GTACAGAGGTGGTGGTCCTCCAGGAACGTGG 3’

*kcnh2a*_exon_5_rev 5’ GTCTGTGGTGAATACACCATGTGTGCCTTTTAAAT 3’

*kcnh2a*_exon_6_for 5’ CAGCAGCATTTACTTCAATCACGATGATCAC 3’

*kcnh2a*_exon_6_rev 5’ GGTTTGGAGCCACCTGAGTGTGAG 3’

*kcnh2a*_exon_7_for 5’ GCACCCTAGCAACTGTAATAACTGCTGC 3’

*kcnh2a*_exon_7_rev 5’ GAATAAGCACTTAGCAGCACTAACCTCCG 3’

*kcnh2a*_exon_8-9_for 5’ GTTCAACCACTGGGTGTCAAACTTACACACT 3’

kcnh2a_exon_8-9_rev 5’ GGGTGGGGAGTGTCTGGGTAAGGG 3’

*kcnh2a*_exon_10-12_for 5’ GGCATGCAGCGCCACCTGCTGTTGATTTC 3’

*kcnh2a*_exon_10-12_rev 5’ GGTACGGCAACTATTCTCCCAATCAACATAC 3’

*kcnh2a*_exon_13-14_for 5’ GAGGTTAGAATGACTTGAATGAGTATAATCAGTG 3’

*kcnh2a*_exon_13-14_rev 5’ GACCCAGCCGAGGCTTGAACCA 3’

*kcnh2a*_exon_15_for 5’ GAGGGTGTGTGATATCAATAGCATGGGG 3’

*kcnh2a*_exon_15_rev 5’ GCGTTCACACAGCACAGCGTAACCAG 3’

*runx2b*_forward 5’ AGCTTCACCCTGACGATTACA 3’

*runx2b*_reverse 5’ CCAGTTCACTGAGACGGTCA 3’

*rpl8*_forward 5’ CCGAGACCAAGAAATCCAGA 3’

*rpl8*_reverse 5’ GAGGCCAGCAGTTTCTCTTG 3’

## ACKNOWLEDGEMENTS

We thank the University of Oregon Aquatic Animal Care Services for animal husbandry; A. Lasseigne and A. Miller for assisting with blastula transplantations; J. Chehab for sequence analysis; H. Markovic for assistance with inhibitor studies; the Stankunas lab for discussions; C. Kimmel, D. Grimes, and V. Lewis for manuscript feedback; and K. Poss for providing the *Tg(tph1b:mCherry)* line.

## Author contributions

S. S., H. K. L., G. A. Y., and K. S. designed experiments. S. S., H. K. L., G. A. Y., A. L. H., A. E. R., and J. A. B. performed experiments. S. S. and K. S. prepared and wrote the manuscript with input from all authors.

## Data and material availability

RNA-Seq data are deposited at the NCBI Gene Expression Omnibus GSE137352. Requests for materials should be addressed to S. S. and K. S.

