## Supplementary Information for "*longfin* causes *cis*-ectopic expression of the *kcnh2a ether-a-go-go* K^+^ channel to autonomously prolong fin outgrowth"

**Figures S1-S7.**

**Table S1.**

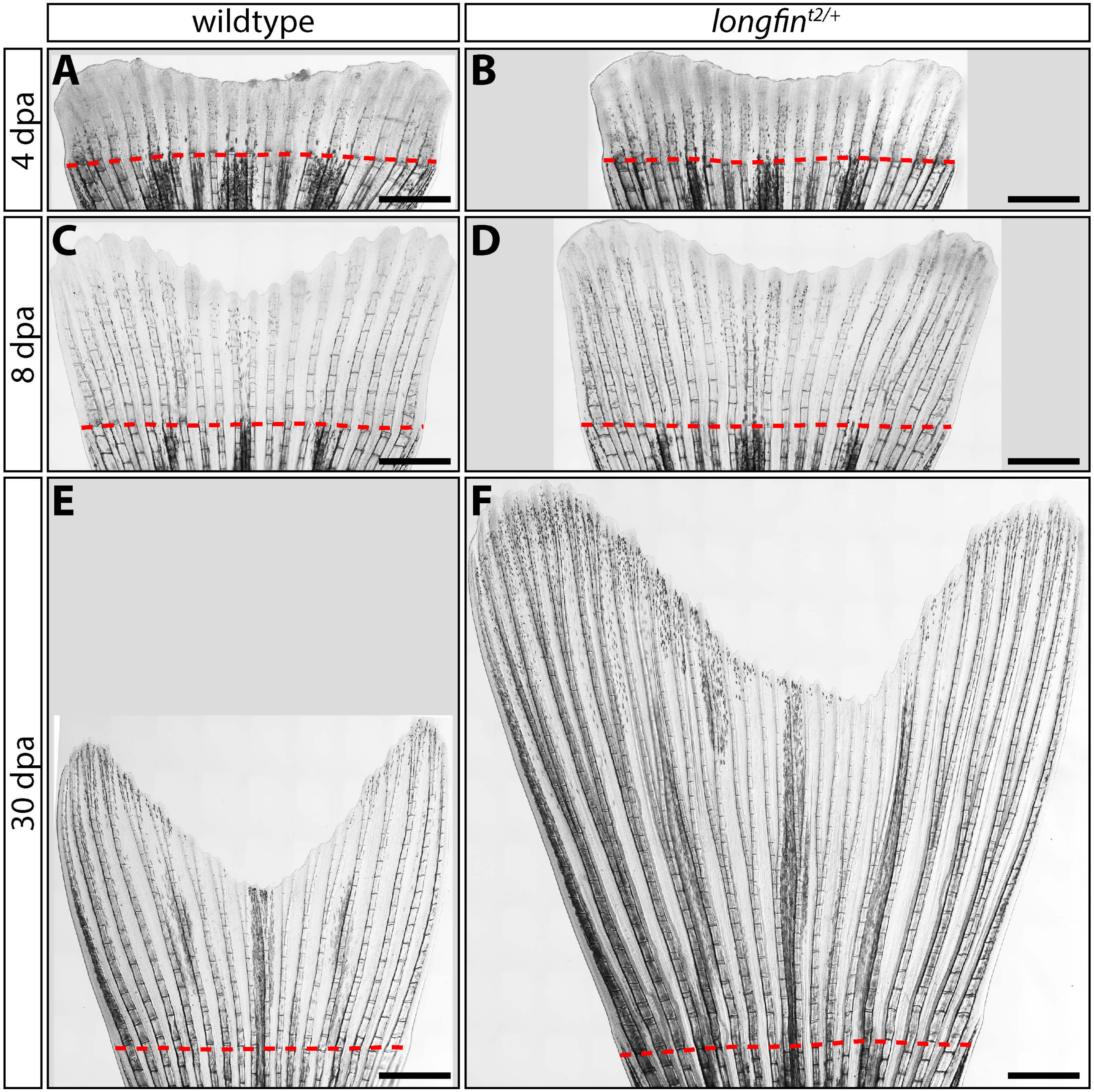


**Supplemental Figure 1**

**Figure S1. Regenerative outgrowth of *lof*$^{t2/+}$ caudal fins fails to decelerate.**

**(A-F)** Stitched differential interference contrast (DIC) microscope images of caudal fins from clutchmate wildtype and *lof*$^{t2/+}$ animals at the indicated day post amputation (dpa). Outgrowth is similar through 8 dpa but persists in *lof*$^{t2/+}$ fish, leading to exceptionally long fins by 30 dpa. Quantitative data for the entire time course (n $\geq$ 12) is shown in Fig. 3. Dashed red lines highlight amputation planes. Scale bars represent 1 mm.

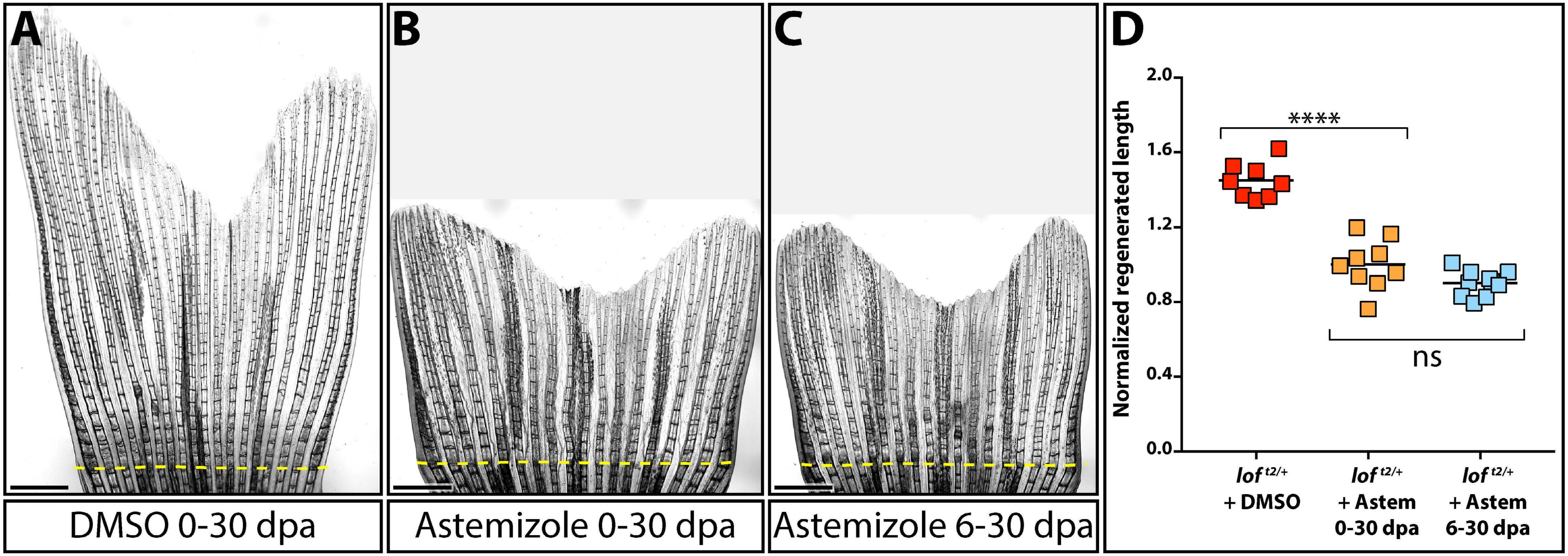


Supplemental Figure 2

**Figure S2. Kcnh2a inhibition during late stage regeneration is sufficient to suppress *lof*$^{t2}$ overgrowth.**

**(A-C)** Representative DIC images showing 30 day outgrowth of *lof*$^{t2/+}$ fins after treatment with (A) DMSO, (B) astemizole (500 nM) from 0-30 dpa, and (C) astemizole (500 nM) from 6-30 dpa. The scale bar indicates 1 mm and the amputation plane is shown by a dashed yellow line. **(D)** Graph showing normalized growth of ray 3 measured from the amputation site; each data point represents an individual animal. Either astemizole regimen blocked overgrowth compared to DMSO ($p < 0.05$) but were themselves statistically indistinguishable assessed by a one-way ANOVA and Tukey's multiple comparison tests.

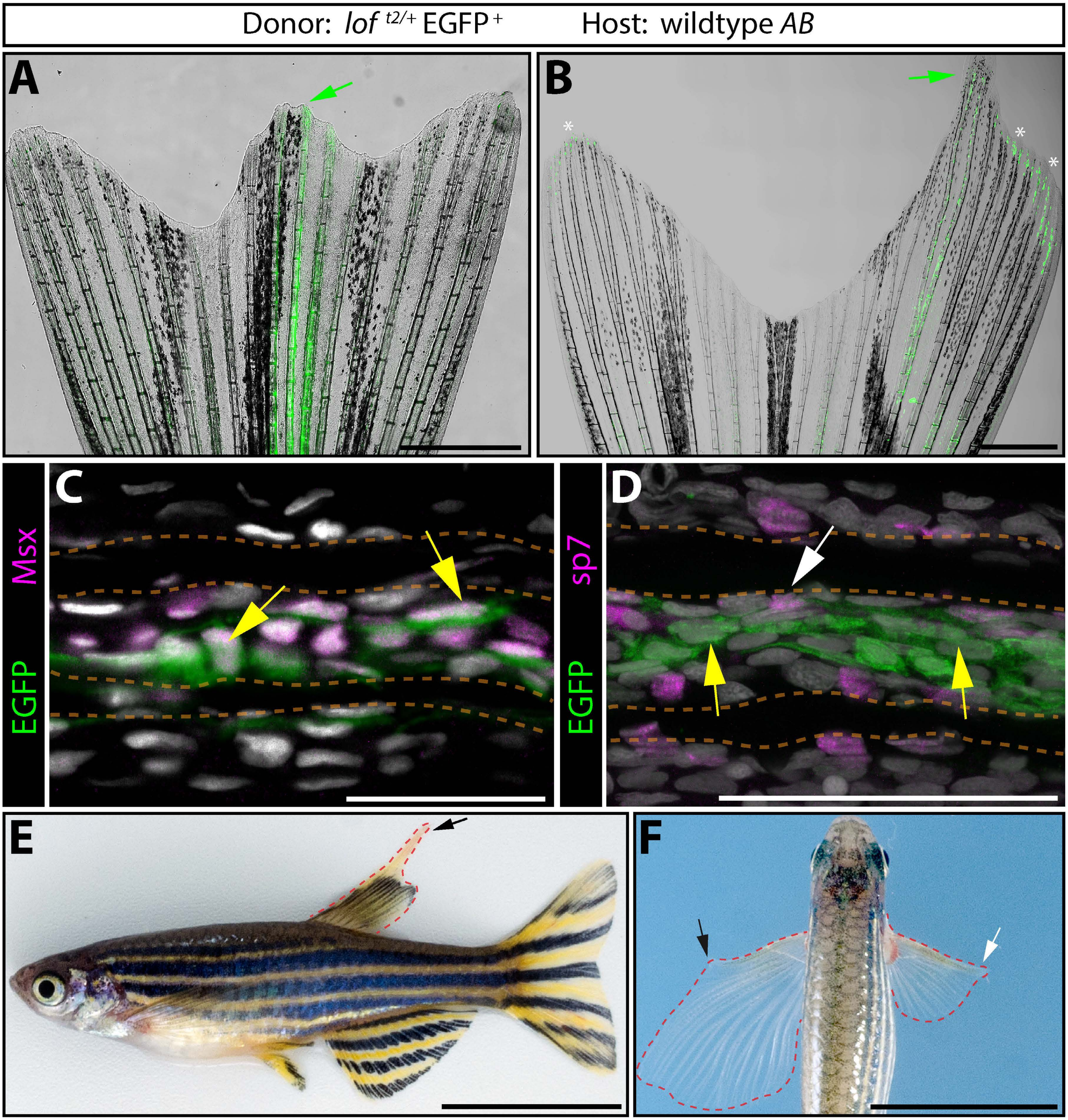


**Supplemental Figure 3**

**Figure S3. *lof*$^{t2}$-*AB* chimeras reinforce *lof*$^{t2}$-ectopic *kcnh2a* acts within the intra-ray mesenchyme lineage.**

**(A, B)** Superimposed epifluorescence/DIC images showing two additional examples of overgrown caudal fin rays from chimeric animals generated by transplanting blastula stage EGFP$^{+}$ *lof*$^{t2/+}$ cells into wildtype *AB* embryos. Green arrows indicate overgrown rays containing EGFP$^{+}$ cells derived from *lof*$^{t2/+}$ embryos and white asterisks point to auto-fluorescent pigment cells. Scale bars indicate 1 mm. **(C, D)** Immunostaining using EGFP and (C) Msx or (D) sp7 antibodies on sections from overgrown chimeric caudal fins generated by blastula-stage transplantation (upper panel). In (C), yellow arrows indicate EGFP$^{+}$/Msx$^{+}$ intra-ray mesenchymal cells derived from *lof*$^{t2/+}$ transplants. In (D), the yellow arrows indicate the EGFP$^{+}$ mesenchyme, the white arrow points to a sp7$^{+}$ osteoblast. Dashed orange lines outline fin ray bones. Scale bars represent 50 μm. **(E)** Lateral whole animal view of a distinctive *lof*$^{t2}$-*AB* chimera. The black arrow indicates overgrown dorsal fin tissue. The scale bar represents 1 cm. **(F)** Dorsal view of a *lof*$^{t2}$-*AB* chimera to highlight contralaterally distinct pectoral fin sizes. Dashed red lines outline the pectoral fins. The black and white arrows point to overgrown and normal sized fins, respectively. The scale bar represents 1 cm.

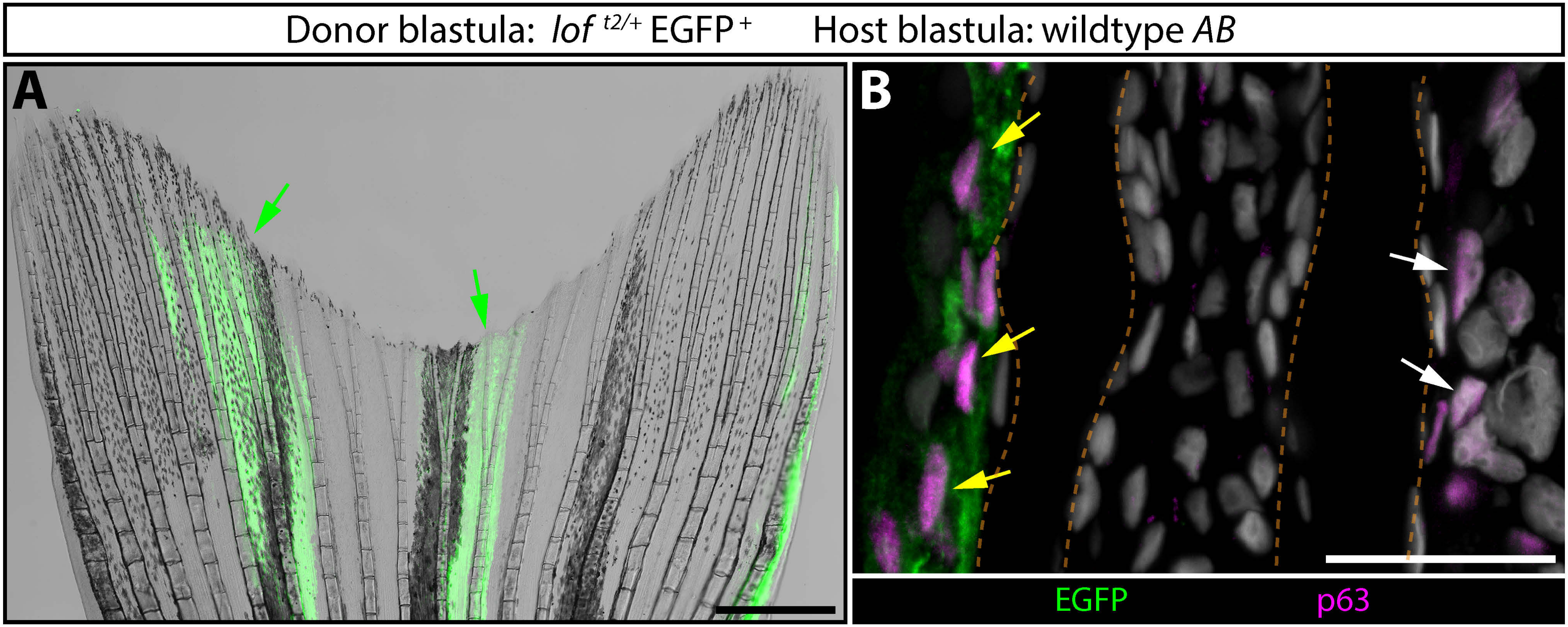


**Supplemental Figure 4**

**Figure S4. Epidermal *lof*$^{t2}$-AB chimeric caudal fins do not overgrow.**

**(A)** A DIC/fluorescence-imaged caudal fin from a *lof*$^{t2}$*-AB* chimera with EGFP$^{+}$ epidermis (green arrows). The scale bar represents 1 mm. **(B)** Immunostaining of sections from the same fin using EGFP and p63 antibodies to detect transplanted *lof*$^{t2/+}$ cells (green cells) and epidermal cells (magenta nuclei), respectively. Yellow arrows point to chimeric EGFP$^{+}$/p63$^{+}$ epidermal cells and white arrows indicate p63$^{+}$ host-derived epidermis. Nuclei are Hoechst-stained (gray), dashed orange lines outline fin rays, and the scale bar shows 50 μm.

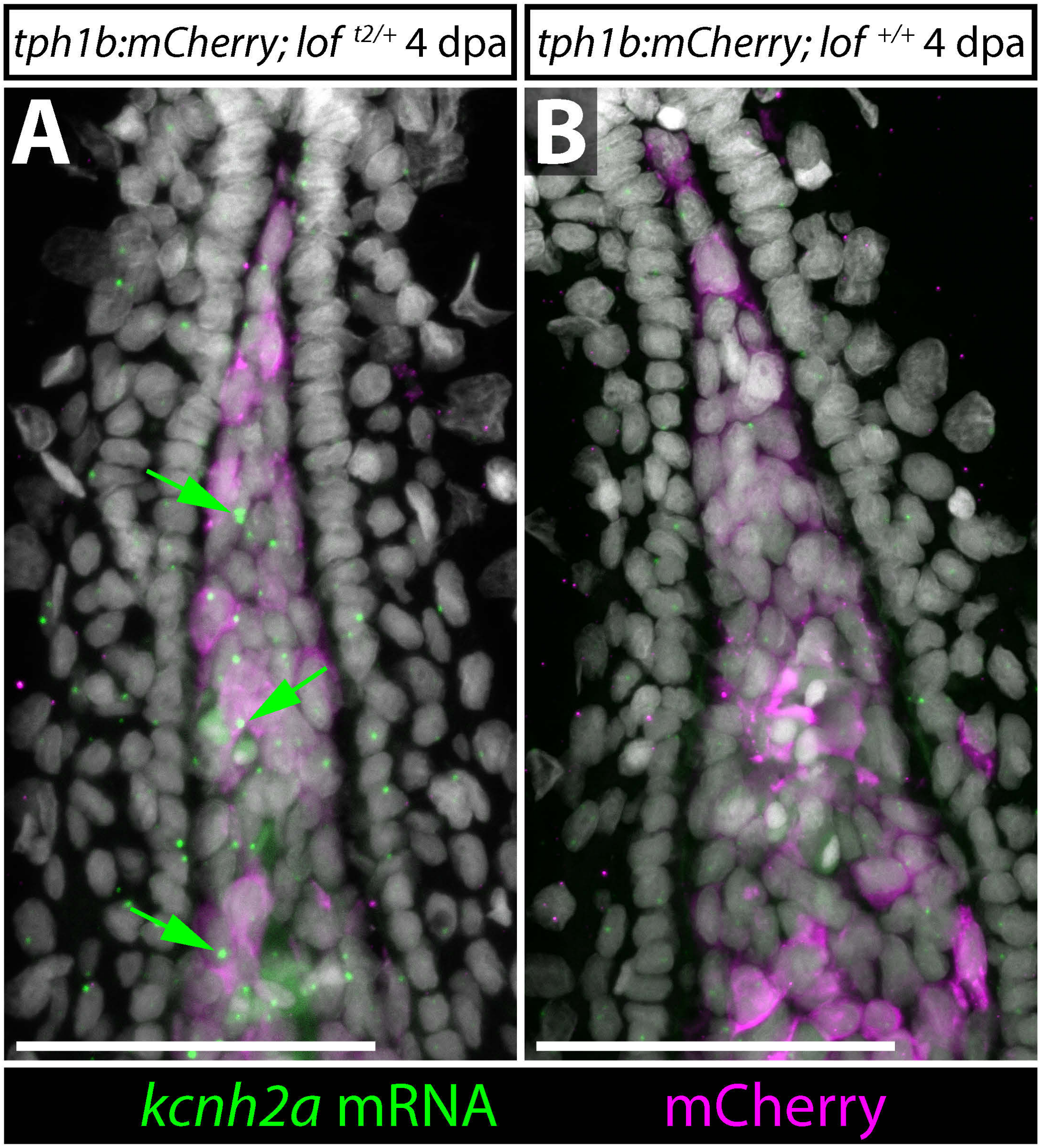


**Supplemental Figure 5**

**Figure S5. *kcnh2a* is expressed in *tph1b:mCherry*-positive distal blastema cells of *lof*$^{t2}$ regenerating caudal fins.**

**(A, B)** Confocal maximum intensity projection images showing combined *kcnh2a* RNAscope and mCherry antibody staining on 4 dpa fin sections from (A) *tph1b:mCherry; lof*$^{t2/+}$ and clutchmate (B) *tph1b:mCherry; lof*$^{+/+}$ animals. Green arrows point to *tph1b:mCherry; lof*$^{t2/+}$ distal blastema cells expressing *kcnh2a* mRNA. Hoechst-stained nuclei are in gray. The scale bars represent 50 μm.

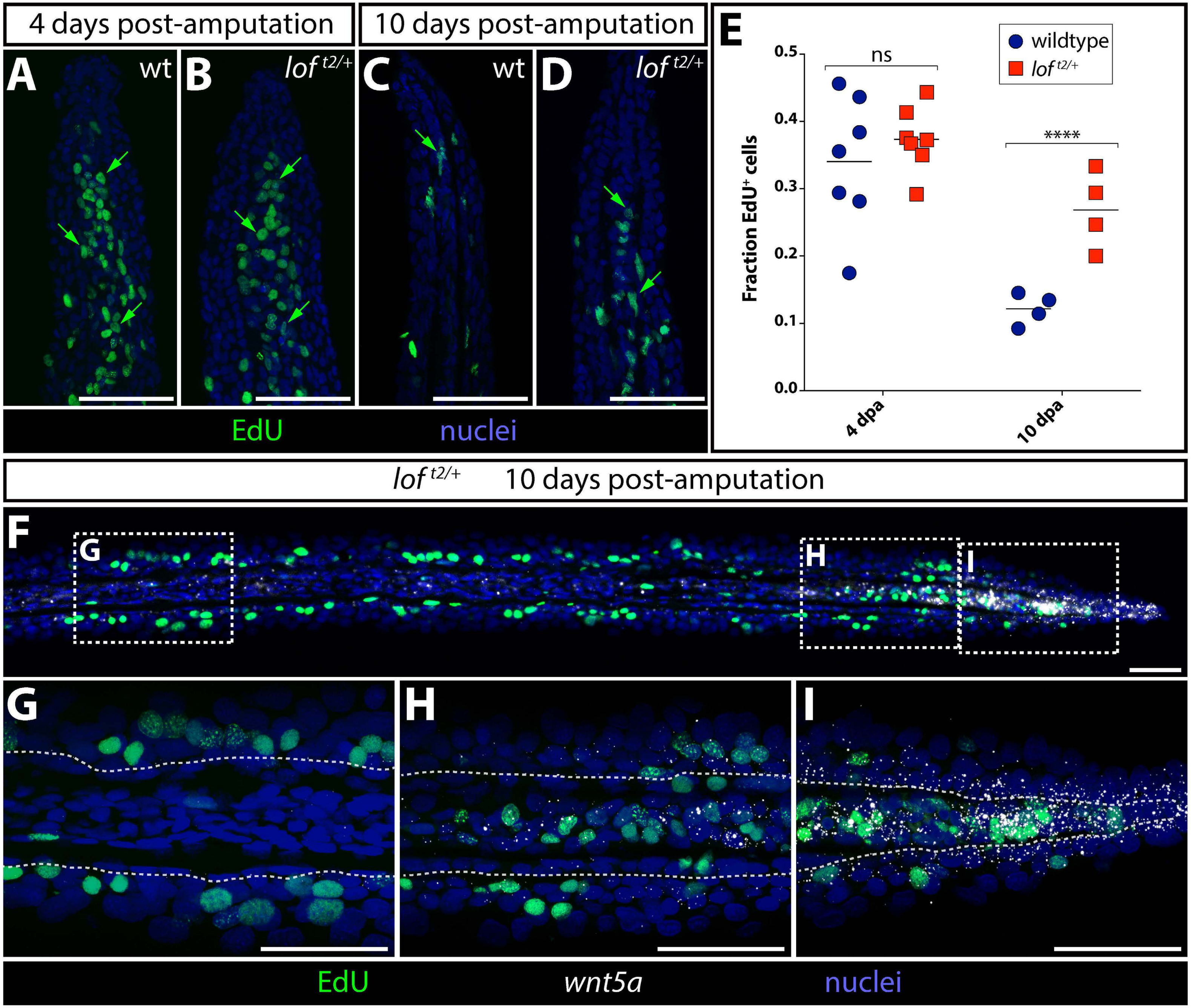


**Supplemental Figure 6**

**Figure S6. *lof*$^{t2}$ regenerating fins have a prolonged cell proliferation period rather than elevated maximal rate**. **(A-D)** Representative confocal images of sectioned 4 and 10 day post amputation regenerating fins showing EdU labeling (green) in wildtype (A, C) and *lof*$^{t2/+}$ (B, D) distal fin mesenchyme (green arrows). Hoechst-stained nuclei are blue. Scale bars are 50 μm. **(E)** Graph showing the fraction of EdU-incorporating distal mesenchymal cells at each time point and between genotypes. Data points represent individual animals. Asterisks indicate $p < 0.01$ by two-way ANOVA with Tukey's multiple comparison tests. ns: not significant. **(F-I)** Confocal images showing the distribution of EdU-incorporating cells (green) and RNAscope for the distal blastema growth factor *wnt5a* (white) on a 10 dpa *lof*$^{t2/+}$ fin section. Dashed boxes in (F) show regions shown at higher magnification in (G-I). Nuclei are Hoechst-stained (blue). Scale bars represent 50 μm.

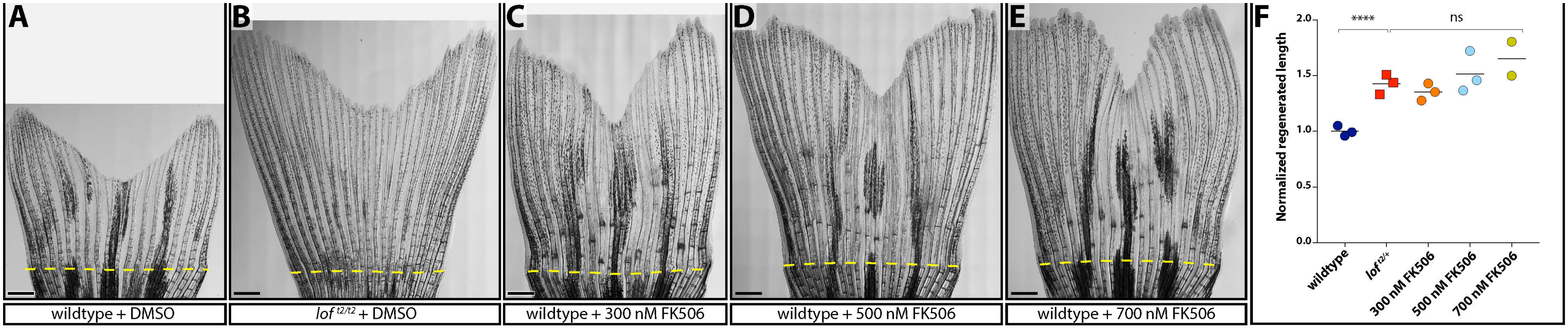


**Supplemental Figure 7**

**Figure S7. Daily acute treatment with FK506 is sufficient to cause fin overgrowth. (A-E)** Representative stitched DIC images showing control DMSO-treated wildtype (A) and *lof*$^{t2/t2}$ (B) caudal fins at 23 dpa. Examples of caudal fins from three cohorts treated for 4 hours daily with 300 (C), 500 (D), or 700 (E) nM FK506 from 2-23 dpa. A dashed yellow line indicates the amputation site. The scale bar represents 1 mm. (F) Quantification of lengths of ray 3 from the amputation site to the end of the fin were measured and normalized to wildtype DMSO-treated samples. Each point represents an individual animal. Shown are DMSO-treated wildtype and *lof*$^{t2/+}$ animals (dark blue circles and red squares, respectively), and wildtype fish treated with the indicated doses of FK506 (orange circles 300 nM, light blue circles 500 nM, gold circles 700 nM). Asterisks: $p < 0.001$ using a one-way ANOVA and a Tukey's multiple comparison test; ns: not significant.

**Table S1. RNA-Seq table showing differentially expressed transcripts in *lof*$^{t2}$ vs. wildtype fins at 4 dpa.**
